# KDM6B-dependent chromatin remodelling underpins effective virus-specific CD8^+^ T cell differentiation

**DOI:** 10.1101/2020.02.03.933218

**Authors:** Jasmine Li, Kristine Hardy, Moshe Olshansky, Adele Barugahare, Linden J. Gearing, Julia E. Prier, Xavier Y.X. Sng, Michelle Ly Thai Nguyen, Dana Piovesan, Brendan Russ, Nicole L. La Gruta, Paul J. Hertzog, Sudha Rao, Stephen J. Turner

**Affiliations:** Department of Microbiology, Biomedicine Discovery Institute, Monash University, Clayton, VIC 3800, Australia; Epigenetics and Transcription Laboratory Melanie Swan Memorial Translational Centre, Sci-Tech, University of Canberra, Bruce 2617 ACT, Australia; Computational Biology & Bioinformatics, Baker Heart & Diabetes Institute, Melbourne, VIC 3004, Australia; Hudson Institute for Medical Research, Clayton, VIC 3168, Australia; Department of Microbiology and Immunology, the Doherty Institute, University of Melbourne, Parkville VIC 3010, Australia; Department of Biochemistry and Molecular Biology, Biomedicine Discovery Institute, Monash University, Clayton, VIC 3800, Australia; Department of Microbiology and Immunology, University of California, San Francisco, California, U.S.A.; QIMR Berghofer Gene Regulation and Translational Medicine laboratory, Department of Immunology, QIMR Berghofer Medical Research Institute, Brisbane, QLD, Australia

**Keywords:** CD8^+^ T cell, epigenetics, histone demethylase, T cell memory, virus immunity, T cell activation

## Abstract

Naive CD8^+^ T cell activation results in an autonomous program of cellular proliferation and differentiation. However, the mechanisms that underpin this process are unclear. Here we profiled genome-wide changes in chromatin accessibility, gene transcription and the deposition of a key chromatin modification (H3K27me3) early after naive CD8^+^ T cell activation. Rapid upregulation of the histone demethylase, KDM6B, prior to first cell division was required for initiating H3K27me3 removal at genes essential for subsequent T cell differentiation and proliferation. Inhibition of KDM6B-dependent H3K27me3 demethylation limited the magnitude of an effective primary virus-specific CD8^+^ T cell response and the formation of memory CD8^+^ T cell populations. Accordingly, we define the early spatio-temporal events underpinning early lineage-specific epigenetic reprogramming that is necessary for autonomous CD8^+^ T cell proliferation and differentiation.

## INTRODUCTION

Upon virus infection, naïve, cytotoxic T lymphocyte (CTL) activation results in a largely autonomous program of differentiation that results in proliferation and acquisition of lineage-specific effector functions (van Stipdonk et al., 2003). The acquisition of lineage-specific CTL functions, such as production of pro-inflammatory cytokines interferon (IFN)-γ, tumour necrosis factor (TNF) (Denton et al., 2011; La Gruta et al., 2004), and expression of cytolytic effector molecules (Jenkins et al., 2007; Kagi et al., 1994; Moffat et al., 2009; Peixoto et al., 2007) helps limit and clear virus infection. Once the infection is cleared, the expanded effector CTL population contracts, leaving a pool of long-lived, pathogen-specific memory T cells. In contrast to naïve CD8^+^ T cells, virus-specific memory CTLs are able to respond more readily and rapidly to subsequent infections without the need for further differentiation (Kaech et al., 2002; La Gruta et al., 2004; Lalvani et al., 1997; Oehen and Brduscha-Riem, 1998; Veiga-Fernandes et al., 2000). This function enables rapid control of a secondary infection leading to immune protection.

Optimal virus-specific CD8^+^ T cell differentiation is underpinned by the coordinated expression of several transcription factors (TFs). The BATF TF has been shown to act early after activation and works in tandem with IRF4 and JUN family members to regulate transcriptional activation of gene loci involved in early immune T cell activation, cell survival and metabolic pathways (Kurachi et al., 2014; Xin et al., 2016). Failure to engage BATF/JUN/IRF4 dependent programs results in diminished CD8^+^ T cell expansion and function. BATF/JUN/IRF4 activity also results in subsequent upregulation of other TFs such as T-BET (encoded by *Tbx21)*, RUNX3 and BLIMP1 (encoded by *Prdm1*), all known to be essential for effective CD8^+^ T cell differentiation (Cruz-Guilloty et al., 2009; Kallies et al., 2009; Kurachi et al., 2014; Wang et al., 2018; Xin et al., 2016). The activation of T-BET and RUNX3 consolidate commitment to the effector CTL lineage (Cruz-Guilloty et al., 2009; Intlekofer et al., 2008; Intlekofer et al., 2005), whilst BLIMP1 is required for terminal effector CTL differentiation (Kallies et al., 2009). These data demonstrate that the stepwise progression of TF expression during virus-specific CTL differentiation is critical for optimal responses. Interestingly, while at little as 2 hours (hrs) of antigenic stimulation is sufficient to initiate CD8^+^ T cell proliferation (van Stipdonk et al., 2001), sustained stimulation for at least 20 hrs is required to install an optimal effector response (van Stipdonk et al., 2003). While this suggests that extensive cellular reprogramming prior to the first cell division is required to ensure optimal CD8+ T cell responses, the exact molecular events that trigger this process are not fully understood.

Within eukaryotic cells, DNA is wrapped around a complex of histone proteins known as a nucleosome, with the nucleosome-DNA complex termed chromatin. Post-translational modification (PTM) of histones is an important mechanism for directing gene expression programs necessary for the process of differentiation in an array of cellular contexts. Histone PTMs contribute to regulation of transcription by providing a platform that promotes binding of TFs and chromatin remodelling proteins (Kouzarides, 2007). We and others, have demonstrated that virus-specific CTL differentiation is associated with genome wide changes in chromatin accessibility and PTMs (Denton et al., 2011; Northrop et al., 2008; Russ et al., 2014; Russ et al., 2017; Scott-Browne et al., 2016; Sen et al., 2016; Wang et al., 2018; Wei et al., 2009; Zediak et al., 2011). More recently, extensive changes in chromatin accessibility, indicative of transcriptional activation was shown to occur prior to first cell division, and was dependent on RUNX3 (Wang et al., 2018). Our own analysis showed that in the naive state, there is co-deposition of histone modifications associated with transcriptional activation (H3K4me3) and repression (H3K27me3) at CD8^+^ T cell lineage specific gene promoters and enhancers (Russ et al., 2014; Russ et al., 2017). Upon T cell activation, loss of H3K27me3 at gene promoters and enhancers was broadly associated with transcriptional upregulation of these poised genes (Russ et al., 2014). These data suggest that the presence of H3K4me3 at specific gene loci ensures that the genome of naive CD8^+^ T cells is preconfigured for transcriptional activation, but it is maintained transcriptionally poised via co-localisation of H3K27me3. While removal of H3K27me3 appears to be a key step in the initiation of naive T cell activation, the timing, genomic targets and specific molecular mechanisms of this initiating event remain to be determined.

The removal of H3K27me3 is specifically catalysed by KDM6A and KDM6B demethylases (Agger et al., 2007). During thymic development, H3K37me3 status is a highly dynamic and stage specific, with KDM6A and KDM6B both playing a key role in modulating these patterns (Manna et al., 2015; Zhang et al., 2012). In mature CD4^+^ T cells, KDM6A activity was required for the rapid expression of several key transcription factors, such as T-BET and STAT family members (LaMere et al., 2017). *Kdm6b*-deficient CD4^+^ T cells demonstrate dysregulated and inappropriate fate specification under T_H_ skewing conditions with promotion of T_H_2/T_H_17 lineages at the expense of T_H_1 and FOXP3 T regulatory cells (Li et al., 2014). Together these data suggest that dynamic modulation of H3K27me3 appears to be critical for multiple stages of T cell differentiation, both during development and activation. However, precisely how modulation of H3K27me3 during the very early stages of T cell activation promote effective T cell immunity is not fully understood.

Despite our earlier observations demonstrating that rapid removal of H3K27me3 is a key outcome of naive, virus-specific CD8^+^ T cell activation (Denton et al., 2011; Russ et al., 2014; Russ et al., 2017), there is little understanding of the molecular mechanisms by which removal of H3K27 facilitates CD8 T cell differentiation. Moreover, the respective roles of KDM6A and KDM6B in mediating H3K27me3 removal remain wholly unknown. Here, we determine that KDM6B is rapidly upregulated upon T cell activation and prior to first cell division. This coincides with a step-wise engagement of transcriptional modules that were linked with rapid H3K27me3 demethylation. This occurred at genes involved in a broad range of cellular support processes that underpin optimal T cell activation and proliferation. Initial H3K27me3 demethylation, and increased chromatin accessibility, targeted regions enriched for BATF/IRF/JUN binding sites, with T-BET and GATA TF binding sites (TFBS) evident at later stages of H3K27me3 demethylation. Small molecule and shRNA inhibition of KDM6B-dependent H3K27me3 demethylation limited the magnitude of an effective primary virus specific CD8^+^ T cell response and formation of functional memory CD8^+^ T cell populations capable of recall. Our data show that H3K27me3 methylation acts as a molecular handbrake on the initiation of effective T cell responses, with H3K27me3 demethylation being a key step at the very earliest stages of T cell activation enabling optimal lineage-specific reprogramming of effector and memory CD8^+^ T cells.

## Results

### Rapid upregulation of KDM6b occurs after naive CD8^+^ T cell activation

We have previously demonstrated that specific transcriptional and epigenetic changes occur within 5 hours (hrs) of naive CD8^+^ T cell activation (Denton et al., 2011; Russ et al., 2014; Russ et al., 2017). In particular, we showed that loss of H3K27me3 occurs rapidly after naive T cell activation (Russ et al., 2014). To better understand the global transcriptional changes associated with the rapid loss of H3K27me3, naïve (CD44^int/lo^CD62L^hi^) OT-I CD8^+^ T cells were sorted after *in vitro* activation with their cognate peptide antigen, the ovalbumin (OVA_257-264_, SIINFEKL) N4 peptide. Changes in gene transcription at early (3, 5hrs) and late (24hrs) times post-stimulation were assessed by RNA-seq and differentially expressed genes (DEGs) that were significantly up or down regulated compared to unstimulated cells identified. We initially assessed the transcriptional dynamics of chromatin modifiers at early (3-5hrs) and late (24hrs) time points after activation (**Fig. 1A**). Interestingly, we observed upregulation across the time course of histone methyltransferases, such as *Suv39h1/h2* and *Ezh2/Suz12* that are associated with deposition of H3K9me3 (Rea et al., 2000) and H3K27me3 (Cao et al., 2002), respectively. This is in line with recent reports showing that upregulation of these components are important for optimal effector CD8^+^ T cell responses (Gray et al., 2017; Pace et al., 2018). We also observed that the H3K27me3 demethylase, *Kdm6b*, was transiently upregulated at 3 and 5hrs but returned to levels observed in naive CD8^+^ T cell levels at 24hrs (**Fig. 1A**). *Kdm6b* was the only one of five histone demethylases analysed that was transcriptionally upregulated in response to TCR, while *Kdm6a*, another histone demethylase that is responsible for H3K27me3 removal, was unchanged (**Fig. 1B**).

**Figure 1.**
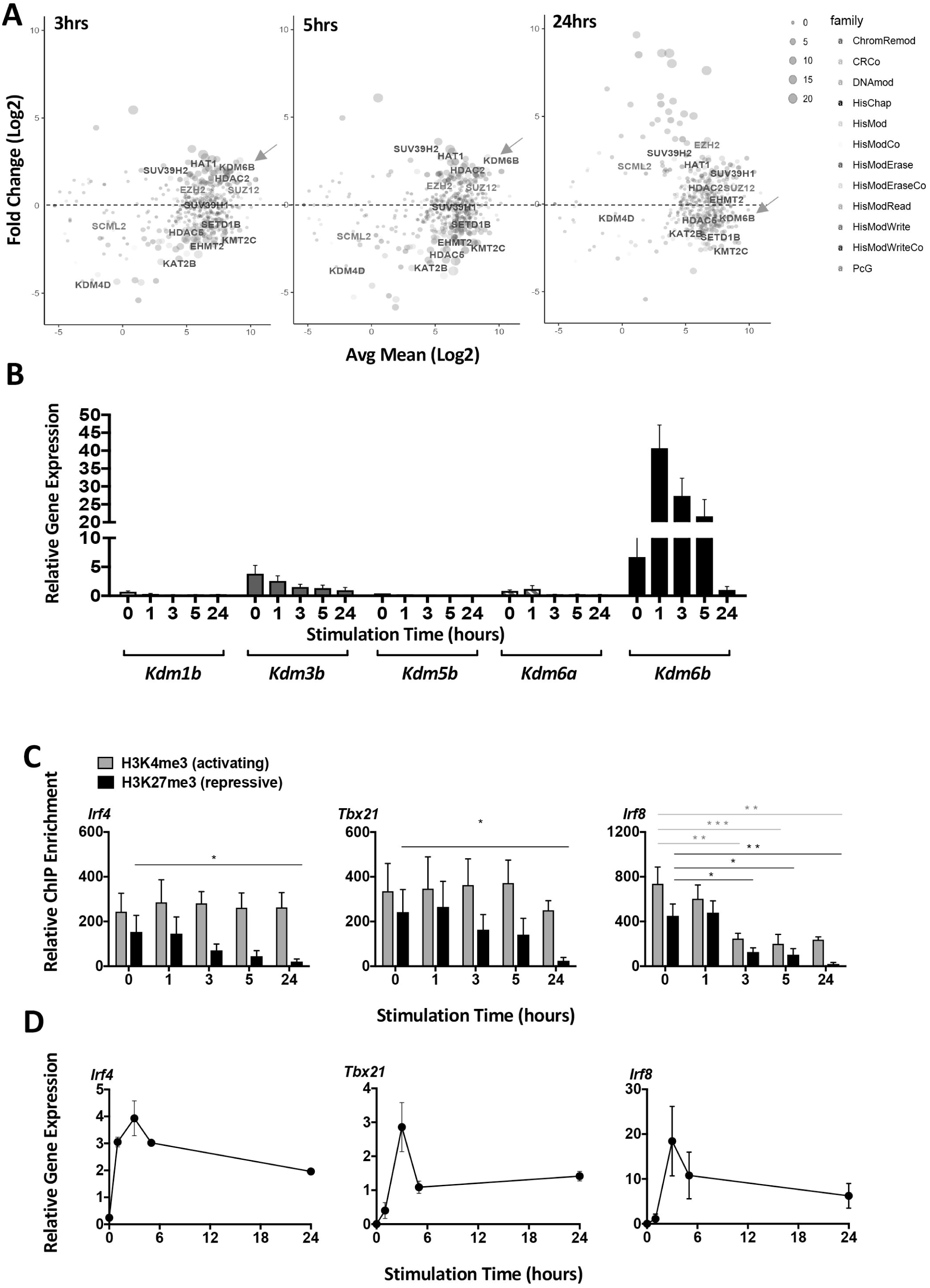
The regulation of Kdm6b and H3K27me3 demethylation during early hours of T cell activation. **A)** Naïve (CD44^lo^CD62L^hi^) CD45.1^+^ CD8^+^ OT-I CTLs were sort purified in and RNA-seq analysis carried on naive CD8^+^ OT-I T cells, or after *in vitro* stimulation with the N4 peptide in the presence of rhIL-2 (10U/mL) for 3, 5 and 24 hours. Expression fold change (log_2_) of histone modifiers by RNA-seq in activated CD8^+^ T cells was compared to unstimulated OT-I naïve CD8^+^ T cells. **B)** Relative gene transcription levels of *Kdm1b, Kdm3b, Kdm5b, Kdm6a* and *Kdm6b* were validated by qPCR. **C)** Relative enrichment of H3K27me3 measured by ChIP-qPCR using primers spanning across the promoter of *Irf4, Tbx21* and *Irf8* gene loci in OT-I CD8^+^ T cells stimulated up to 2hrs in 10U/mL of rhIL-2 with the 1 μg N4 peptide *in vitro*. **D)** Relative gene transcription measured by real time qPCR validated the transcriptional pattern of the transcription factors, *Tbx21* (encodes T-BET), *Irf4* (IRF4) and *Irf8* (IRF8) in OT-I naïve CD8^+^ T cells stimulated as described above. Data are shown as mean ± SEM from 3 independent repeats. Data are shown as mean ± SEM from 3 independent repeats with statistical significance calculated using a one-tailed Student’s T-test (*p<0.05 **p<0.01, ***p<0.001).

### Rapid H3K27me3 demethylation occurs after naive CD8^+^ T cell activation

TCR signalling-induced *Kdm6b* up-regulation coincided with a progressive loss of H3K27me3 at the promoters of the key transcription factors *Irf4, Tbx21* and *Irf8* at 5 and 24hrs after activation, while the levels of H3K4me3 enrichment remained consistent across these time points (**Fig. 1C**). This progressive loss of H3K27me3 appeared to selective as H3K27me3 remained largely constant at the promoter of *MyoD* and *Actin* (**Supplementary Figure S1**). Furthermore, the removal of H3K27me3 coincided with concomitant transcriptional upregulation of these TFs (**Fig. 1D**). These data are consistent with rapid KDM6b dependent H3K27me3 demethylation initiating transcriptional activation after T cell stimulation.

To identify the genomic regions that underwent rapid H3K27me3 demethylation, we carried out H2K27me3 ChIP-seq at 3, 5 and 24hrs after naive T cell activation (**Figure 2**). Consistent with previous reports (Araki et al., 2009; Russ et al., 2014), naive CD8^+^ T cells exhibited broad H3K27me3 regions (**Fig. 2A**). Upon CD8^+^ T cell activation, H3K27me3 domains were trimmed significantly, with this remodelling maintained up to 24hrs (**Fig. 2A)**. H3K27me3 demethylation was evident at 7137, 4022 and 13077 regions at 3, 5 and 24hrs, respectively (**Fig. 2B**), far exceeding the number of regions that had gained H3K27me3 (518, 3641 and 2342 regions, at 3, 5 and 24hrs, respectively) (**Supplementary Figure S2A**). Both gain and loss in H3K27me3 levels occurred directly at the promoter, the transcription start site (TSS), exons, 5’ UTR and 3’ UTR of a gene, with most H3K27me3 changes annotated to introns, intergenic regions and short interspersed elements (SINEs) (**Supplementary Figure S2B**), indicating the role of H3K27me3 in regulating both protein coding and non-coding regions.

**Figure 2.**
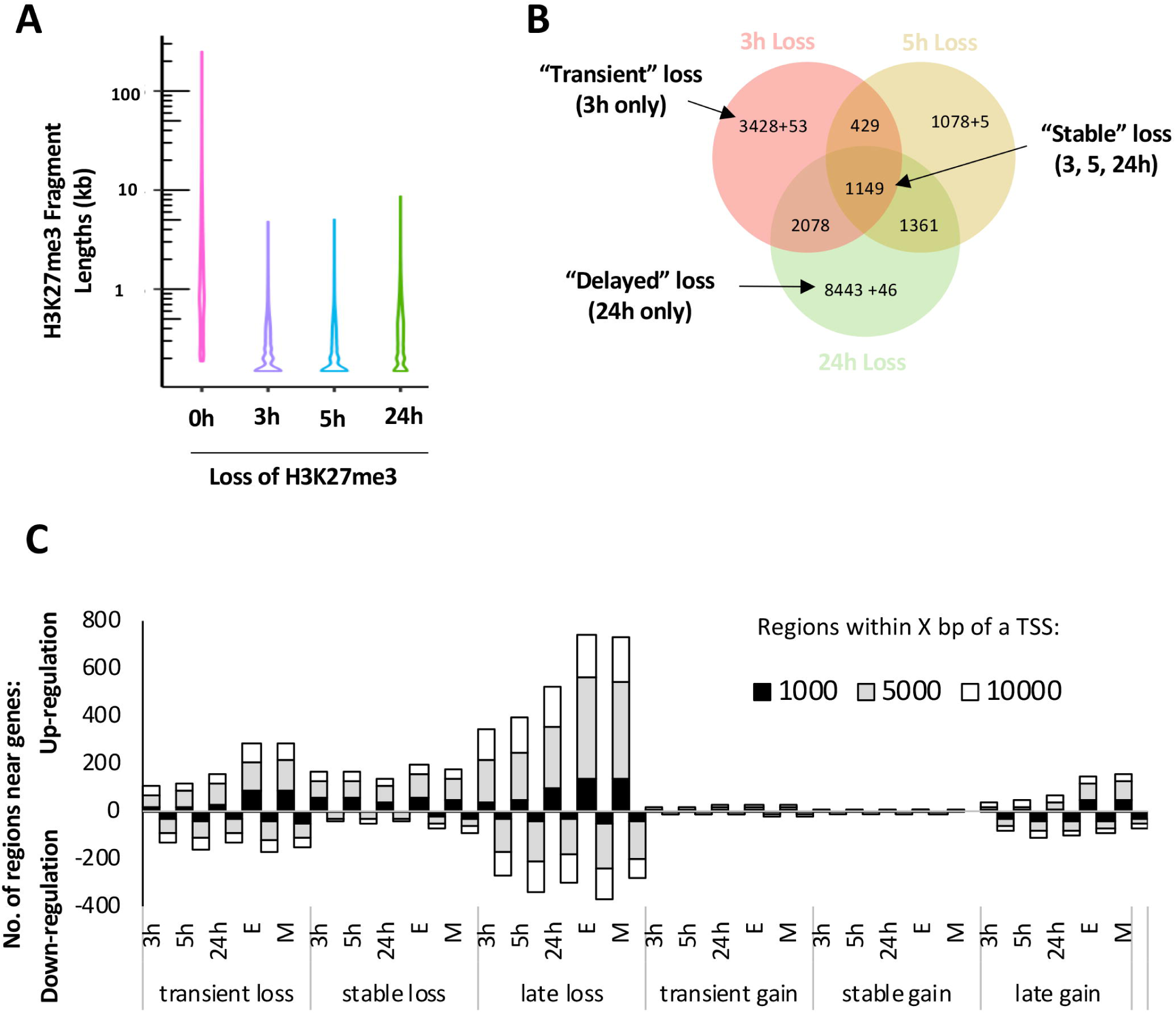
Dynamic regulation of H3K27me3 during early CD8^+^ T cell activation. **A)** H3K27me3 ChIP-seq was performed on either naive CD8^+^ OT-I T cells, or on OT-I T cells after 3, 5, or 24 hrs of *in vitro* stimulation as described above in Figure 1. Data was mapped back to the mouse genome (version mm10). H3K27me3 domain length was assessed within regions that exhibited loss of H3K27me3 within activated CD8^+^ T cells compared to naive CD8^+^ T cells. **B)** Genomic regions that marked with H3K27me3 within OT-I CD8^+^ T cells activated for 3, 5 and 24hrs were enumerated and compared to H3K27me3 regions within unstimulated naive OT-I T cells. Genomic regions exhibiting H3K27me3 loss only after 3hrs of stimulation were characterised as “transient” loss; decreased H3K27me3 at 3, 5 and 24hrs of stimulation were “stable” loss and decreased H3K27me3 at 24hrs only after stimulation as “delayed” loss. **C)** H3K27me3 ChIP-seq was performed on either naive CD8^+^ OT-I T cells, or on OT-I T cells simulated as described above in Figure 1. Data was mapped back to the mouse genome (version mm10). Genomic regions that either lost or gained H3K27me3 within activated CD8^+^ OT-I T cells were compared to the unstimulated sample. These regions were categorised into either “transient”, “delayed” or “stable” loss or gain of H3K27me3. The number of regions in the groups of “transient”, “stable”, “delayed” H3K27me3 loss and gain were within either 1000, 5000 and 10,000 base pairs (bp) of the transcription start site of differentially expressed genes identified after in vitro activation (see Figure 1) or identified after 5 hrs stimulation of *ex vivo*-derived effector OT-I CD8^+^ T cells (Russ et al., 2014).

To investigate the dynamics of H3K27me3 demethylation, we classified regions based on the timing of H3K27me3 removal. We observed that 3428 regions (48%) were transiently demethylated exhibiting a decrease in H3K27me3 at 3hrs (transient loss). In contrast, relatively few regions (1149, 16.1%) were stably demethylated at all time points measured (3, 5 and 24hrs, stable loss). Of the 13077 regions demethylated at 24hrs, the majority of these (8443, 65%) only showed demethylation at the 24hr time point (delayed loss) (**Fig. 2B**). These data are indicative of a staged H3K27me3 demethylation within the first 24hrs of naive T cell activation. Regions with a “transient”, “stable” and “delayed” gain in H3K27me3 were also detected but the numbers of these regions were significantly smaller by comparison (**Supplemental Figure 2A**). Hence, we primarily focused on regions that exhibited H3K27me3 demethylation early after T cell activation.

### Early H3K27me3 demethylation initiates cellular processes required during effector and memory differentiation

To understand how early H3K27me3 demethylation impacted the gene expression profiles observed in early activated CD8^+^ T cells, H3K27me3 demethylated regions were annotated to nearest neighbouring DEGs (±10kb). Within early hours of T cell activation, the majority of the regions exhibiting stable and delayed demethylation were associated with transcriptionally active genes, rather than down-regulated genes (**Figure 2C**). Importantly, this association continued to be evident in *ex vivo*-derived effector and memory CD8^+^ OT-Is (**Figure 2C**) indicating that these early changes in H3K27me3 demethylation are associated with transcriptional upregulation and are transmitted into effector and memory CD8^+^ T cells.

To better visualise the changes in H3K27me3 deposition for the genes that were differentially regulated after activation, we generated heat maps showing the tag density within H3K27me3 peaks (± 5kb from the centre of the peak) annotated to the nearest neighbouring gene (**Fig. 3A**). Interestingly, regions with that exhibited rapid demethylation that was stable at the 24hr time point were linked to genes with a well-documented role in T cell biology, such as *Irf4, Irf8, Tbx21, Il10, Zeb2, Prdm1, Atf3, Lag3, Cd83, Ccl1* and cell cycle processes such as *Nek8 and Cdkn1a* (**Fig. 3A**).

**Figure 3.**
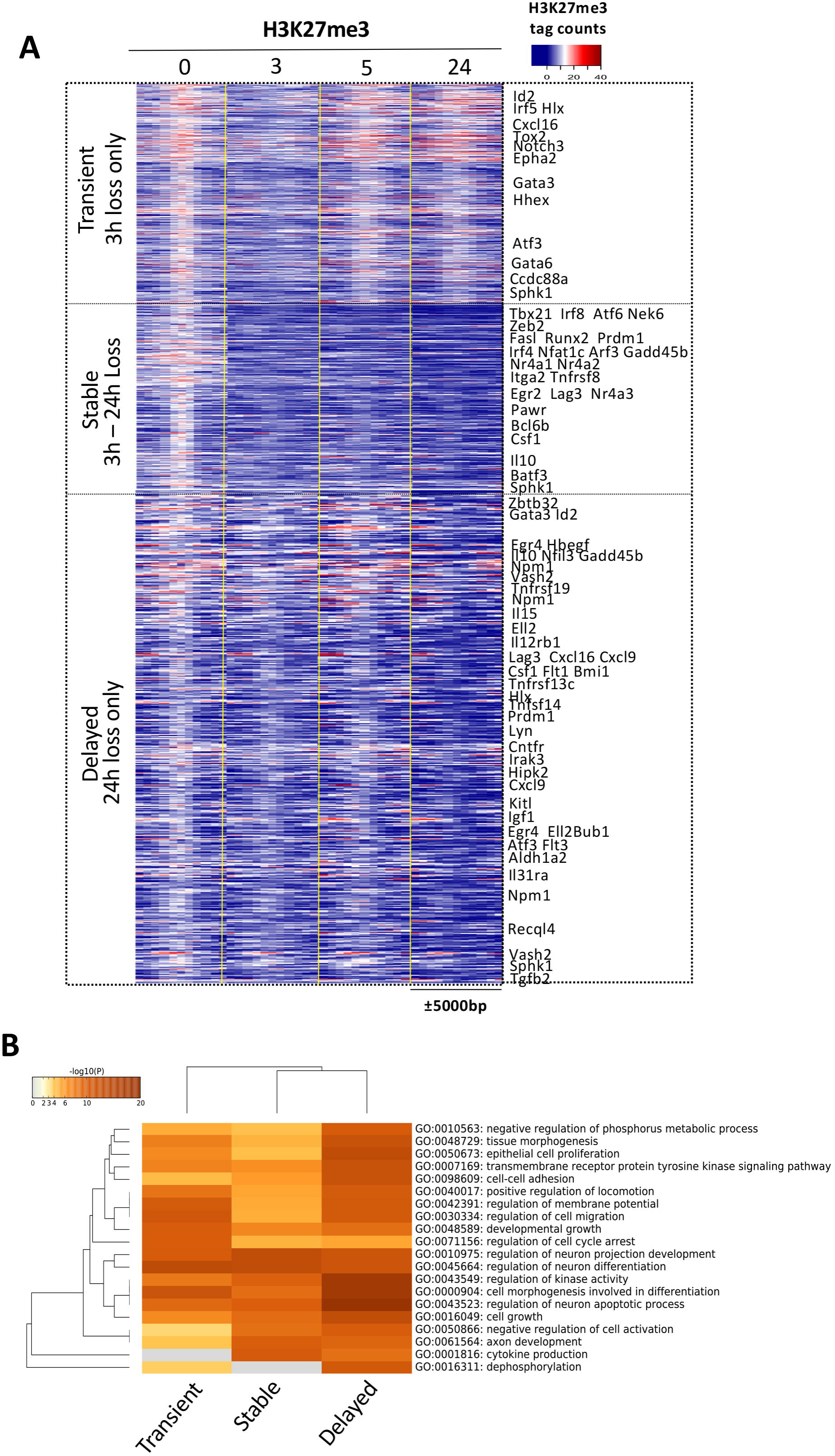
H3K27me3 demethylation regulates genes involved in cellular support processes. **A)** The total number of H3K27me3 sequence tags within ± 5 kb of the middle of the peak was transformed (log2) and converted into a heatmap according to (Russ et al., 2014). Hierarchical clustering was then used to identify genomic regions that exhibited similar patterns of transient, stable or delayed H3K27me3 loss. These regions were then annotated to nearest neighbour genes (listed). **B)** Gene Ontology (GO) analysis of annotated gene loci linked to H3K27me3 regions exhibiting “transient”, “stable” or “delayed” loss of H3K27me3 was carried out and hierarchical clustering based on p value carried out.

To investigate the potential pathways that are impacted by H3K27me3 demethylation upon activation, we carried out gene ontology analysis to identify distinct cellular pathways regulated by distinct H3K27me3 demethylation patterns (**Fig. 3B**). Genes with regions of transient demethylation were primarily involved in chemokine and cellular migration processes. Interestingly, gene loci that exhibited stable and/or delayed demethylation were enriched for general cellular processes such as cell growth, cell morphogenesis, cell adhesion, proliferation and cell cycle arrest (**Fig. 3B**). These data suggest that early H3K27me3 demethylation upon T cell activation is important for the transcriptional activation of cellular support processes required for optimal T cell activation and differentiation.

To explore this further, we carried out gene ontology of DEGs identified at 3, 5 and 24 hrs after activation in our RNA-seq data. K-means clustering partitioned DEGs into modules of transcriptionally-induced (sets a-d) or repressed genes (sets e-h) that exhibited distinct kinetics (**Supplementary Figure S3A, B**). Gene ontology analysis of DEGs that were upregulated after activation showed distinct functional associations depending on when they were upregulated. Genes that were rapidly, but transiently upregulated (set a) were enriched in genes associated with inflammatory and immune response function **(Supplementary Figure S3C**). Genes that were transcribed over the entire time course (sets b, c, 3-24 hrs) were enriched for cellular support processes such as RNA binding and processing, metabolic pathways, and cell cycle/proliferation processes (**Supplementary Figure S3C**). Finally, those genes upregulated at 24hrs only were enriched for DNA repair and cellular division (**Supplementary Figure S3C**). Taken together, our data demonstrate that naive CD8^+^ T cell activation results in rapid H3K27me3 demethylation resulting in step-wise engagement of transcriptional modules important for readying the activated T cell for subsequent proliferation and differentiation.

### H3K27me3 removal establishes a permissive chromatin landscape for transcription factor binding

Having established that H3K27me3 demethylation correlates with transcriptional activation of key cellular processes upon T cell activation, we sought to understand the molecular mechanisms that underlie this process. Formaldehyde-Assisted Isolation of Regulatory Elements (FAIRE)-qPCR and H3 ChIP-qPCR showed T cell activation induced a loss of H3, and concomitant increase in chromatin accessibility at the *Irf4, Tbx21* and *Irf8* promoters (**Fig. 4A**). To assess the link between increased chromatin accessibility and H3K27me3 demethylation at a genome-wide scale, we performed ATAC-seq at early and late activation time points (0, 3 and 24 hrs) and cross-referenced it to the H3K27me3 ChIP-seq data at the same time points. A significantly higher number of regions (27.5%) that exhibited stable H3K27me3 loss at 3hr also exhibited an increase in chromatin accessibility compared to regions with transient or delayed H3K27me3 demethylation (10%). Importantly, regions that exhibited concomitant H3K27me3 loss and increased chromatin accessibility were associated with transcriptionally upregulated genes at the same time points (**Supplementary Figure S4**). Importantly, while genes exhibiting demethylation were upregulated, there were also transcriptional upregulation of genes exhibiting no detectable change in H3K27me3 modification. This may reflect a distinct mechanism for transcriptional regulation.

**Figure 4.**
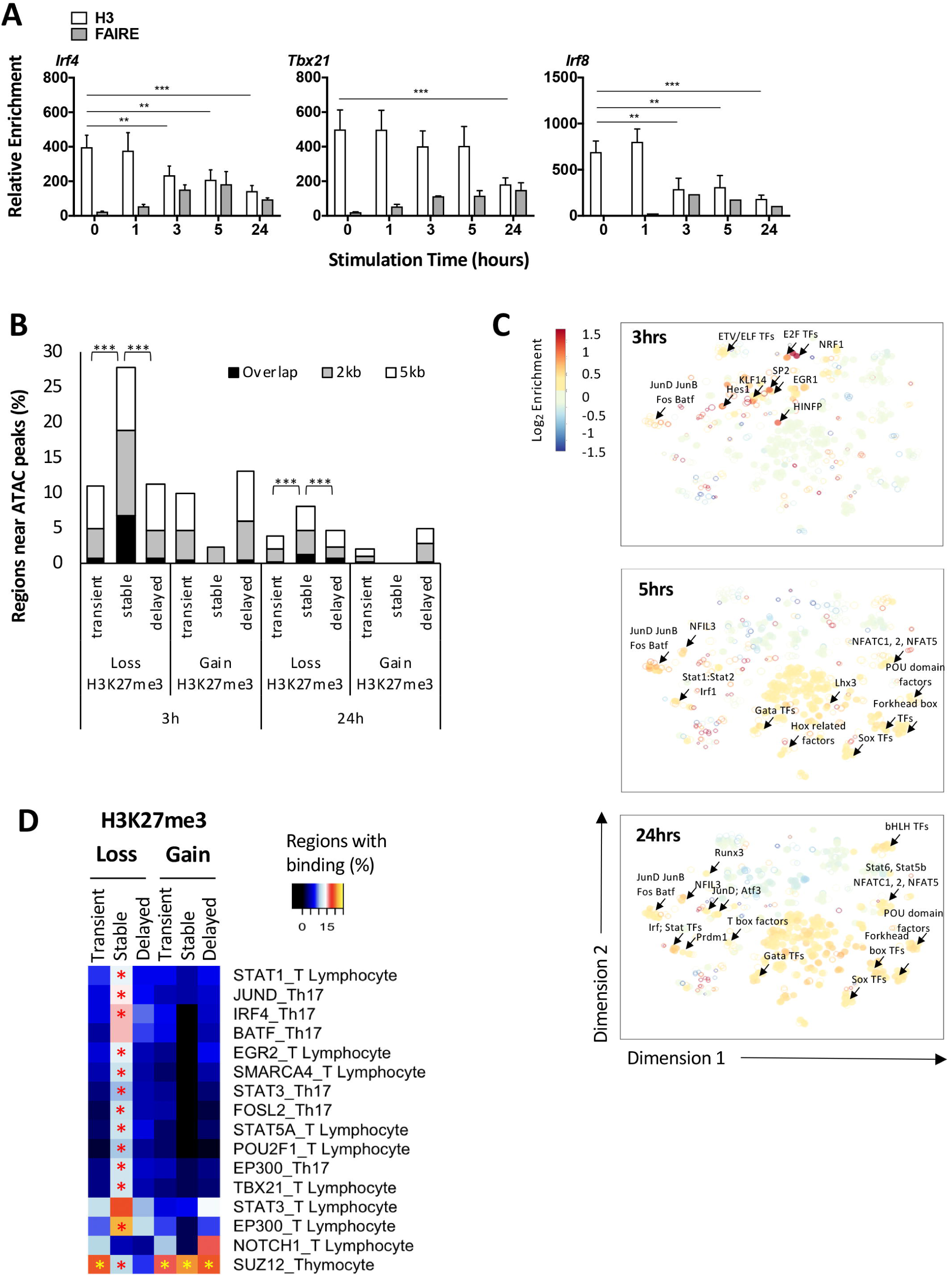
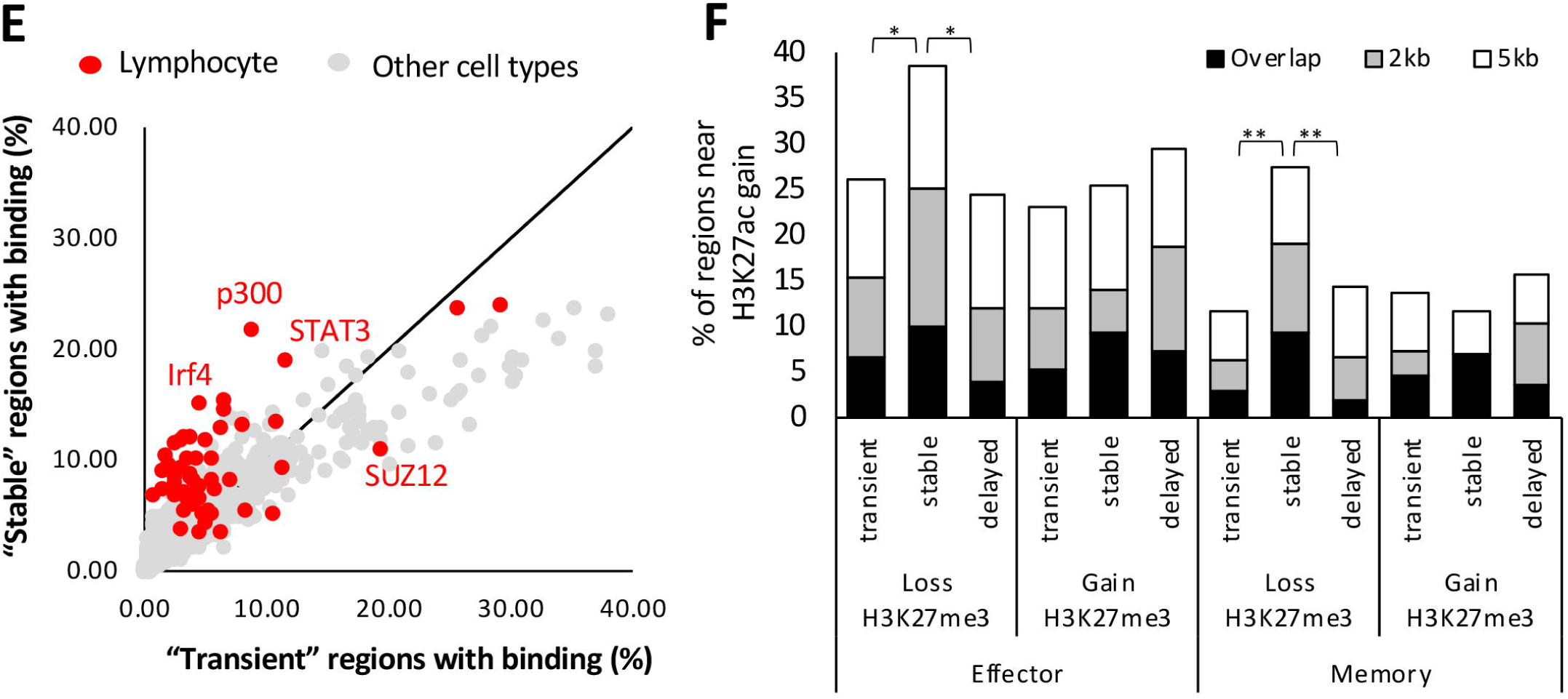
Stable H3K27me3 removal at genomic regions targeted by T cell specific TFs. **A)** Naive OT-I CD8^+^ T cells were activated as previously described above. Changes in chromatin accessibility was assessed by FAIRE-qPCR and H3 histone ChIP using qPCR and primers specific the *Irf4, Tbx21* and *Irf8* promoters. Data are shown as mean ± SEM from 3 independent repeats with statistical significance calculated using a one-tailed Student’s T-test (*p<0.05 **p<0.01, ***p<0.001). **B)** Genome wide changes in chromatin accessibility was assessed by ATAC-Seq on either naive OT-I CD8^+^ T cells, or activated for 3 and 24hrs. Shown is the proportion (%) of regions exhibiting distinct H3K27me3 demethylation dynamics that either directly overlapped or were positioned within 2 to 5kb to the centre of called ATAC-seq peaks. Significantly higher percentage of regions with stable demethylation overlapped or were within <5kb of a chromatin accessible region at 3hr and 24hr than those with transient or delayed demethylation (*** p<2.2e-16). **C)** Prediction of transcription factor binding sites within H3K27me3 demethylated regions was carried out using the CiiiDER algorithm (Gearing et al., 2019; Russ et al., 2017). Shown is a t-SNE plot displaying transcription factor motif enrichment (log_2_) within H3K27me3 demethylated regions at 3, 5 and 24hrs after T cell activation compared to H3K27me3 methylated regions common to all time points (0, 3, 5 and 24hrs). Red circles represent highly enriched TFBS, blue circles represent TFBS that are under-represented. Open circles represent significantly enriched TF motifs that are present in at least 15% of gene loci **D)** Publicly available TF ChIP-seq data for TFs identified to have significantly enriched TFBS (**Fig. 4C**) were downloaded from GEO data sets. The data was mapped to regions exhibiting “transient”, “stable” or “delayed” H3K27me3 demethylation and the percentage (%) of regions exhibiting overlap of TF binding with either H3K27me3 demethylation, or H3K27me3 gain determined. “*” indicates significantly greater binding percentages compared to all H3K27me3 enriched regions at 0hr (p<0.01). **E)** The percentage of genomic regions exhibiting overlap in transcription factor binding in either stable or transient H3K27me3 demethylation were compared. Highlighted in red is publicly available TF ChIP-seq data derived from T cell populations, while TF binding data from other cell types are represented in grey. Gene loci exhibiting significant binding to stable regions or transient regions are named. **F)** The proportion (%) of genomic regions identified to exhibit H3K27me3 demethylation that also overlapped (within 0 to 5kb) to a regions that exhibited H3K27Ac gain within effector or memory OT-I CD8^+^ T cells (Russ et al., 2014) is shown. A significantly higher proportion of regions with stable demethylation overlapped or were within <5kb to a H3K27Ac enriched region in effector (* p<2.2e-16) and memory (** p<6.5e-5) OT-I CD8^+^ T cells than those with transient or delayed demethylation.

A number of these genomic regions exhibiting chromatin remodelling prior to first cell division were also observed in mature *ex vivo* effector and memory IAV-specific CTLs (**Supplementary Figure S4**). These data suggest that early H3K27me3 demethylation upon naive T cell activation results in a profound transition in the chromatin and transcriptional landscape that is maintained in mature virus-specific effector and memory CTLs.

Given the link between loss of H3K27me3 and increased chromatin accessibility, we next assessed which TF binding motifs were enriched at H3K27me3 demethylated regions at 3, 5 and 24 hrs after activation (**Fig. 4C**). Interestingly, in line with the stepwise induction of specific transcriptional modules at distinct stages of early T cell activation (**Supplementary Figure S3**), we observed staged enrichment for specific TF motifs at 3, 5 and 24hrs after T cell activation. ATF/JUN motifs were amongst one of the earliest motifs detected at demethylated regions 3 hours post-activation (**Fig. 4C**). Importantly, the level of ATF/JUN motif enrichment further intensified up to 24hrs. At 5hrs after activation, TF motifs for REL, IRFs/STATs/GATA family members emerged at H3K27 demethylated regions with the appearance of TBX21, RUNX3 and NFIL3 sites at regions demethylated at 24hrs. Interestingly, motifs for EGR and E2F TFs were only transiently enriched (3hrs) and were negatively enriched at regions demethylated at 5 and 24hrs after T cell activation. Hence, dynamic regulation of H3K27 methylation upon naive T cell activation results in ordered chromatin remodelling events that appear to ready the chromatin landscape for specific TF binding.

To determine whether enrichment of TF motifs at H3K27me3 demethylated sites corresponded to specific TF binding, regions that exhibited loss or gain of H3K27me3 were overlaid with publicly available T cell TF ChIP-seq data (**Fig. 4D, E**). Supporting the TF motif enrichment analysis (**Fig. 4C**), stably demethylated regions in recently activated CD8^+^ T cells were enriched for AP-1 (JUND-14.6%, BATF-15.2%, FOSL2-11.6%) and STAT (STAT1-13.3%, STAT3-10.2%, STAT5A-11.8%) members, and TBX21 (12.1%) and IRF4 (15.4%) binding compared to either the transiently or delayed groups which ranged between 2 and 6% in the same dataset (**Fig. 4D, E**). In contrast, 19.3% of the transiently H3K27 demethylated regions and 17-20% of the regions exhibiting H3K27me3 gain overlapped with SUZ12 binding, a component of the PRC2 complex that is responsible for H3K27me3 deposition (**Fig. 4D, E**). This suggests that the observed upregulation of PRC2 components, EZH2 and SUZ12, early after T cell activation may correlate with remethylation of transiently demethylated regions soon after T cell activation.

Importantly, a greater percentage of stably demethylated regions (21.8%) also showed binding for the histone acetyltransferase, p300 compared to the transient demethylated regions (8.8%, **Fig. 4D, E**). The potential for p300 binding at these stably demethylated regions was linked to genes that were transcriptionally induced in effector and/or memory T cells (**Supplementary Figure S4**). These data suggest that p300 binding and subsequent acetylation of H3K27 is required for the demethylation to remain stable instead of transient. To test this, we overlaid our previous H3K27ac data from naïve, effector and memory CD8^+^ T cells (Russ et al., 2017) with transiently, stable and delayed H3K27me3 demethylated regions. In comparison to the transiently demethylated regions, there was a greater proportion of the stably demethylated regions that either overlapped (10%) or were within 2kb (25%) or 5kb (38%) of a region with increased H3K27ac in effector T cells (**Fig. 4F**). This trend was similarly observed in memory T cells albeit at lower percentages (**Fig. 4F**). These overlapping regions with stable demethylation in early hours of T cell activation and effector and memory T cells were positioned near differentially expressed genes linked to H3K27Ac^+^ regions (**Supplementary Figure S4**). Together these data support the notion that early H3K27me3 demethylation enables increased chromatin accessibility and this is further stabilised by the binding of p300 and subsequent H3K27 acetylation. Hence, within the first 24hrs, remodelling of the chromatin landscape provides a platform enabling binding of T cell specific transcription factors. This provides a molecular basis for engagement of the CD8^+^ T cell proliferation and differentiation program induced by T cell activation.

### Inhibition of H3K27 demethylations limits CD8^+^ effector T cell differentiation

To test whether early H3K27me3 demethylation was necessary for induction of appropriate virus-specific T cell differentiation, we treated naive OT-I CD8^+^ T cells with GSK-J4, a small molecule inhibitor, which binds to the catalytic pocket of KDM6B (Kruidenier et al., 2012). This was followed by activation with the N4 peptide for 0, 1, 3, 5 and 24 hours (**Fig. 5A**). GSK-J4 inhibition of KDM6B activity prevented the removal of H3K27me3 across all of the stimulation time points at the *Tbx21, Irf4* and *Irf8* promoters, without affecting pre-existing H3K4me3 levels (**Fig. 5A**). This inhibition of histone demethylase activity significantly reduced the transcriptional levels of *Tbx21, Irf4* and *Irf8* compared to both mock treated, and OT-I T cells treated with a non-functional analogue (GSK-J5) (**Fig. 5B**).

**Figure 5.**
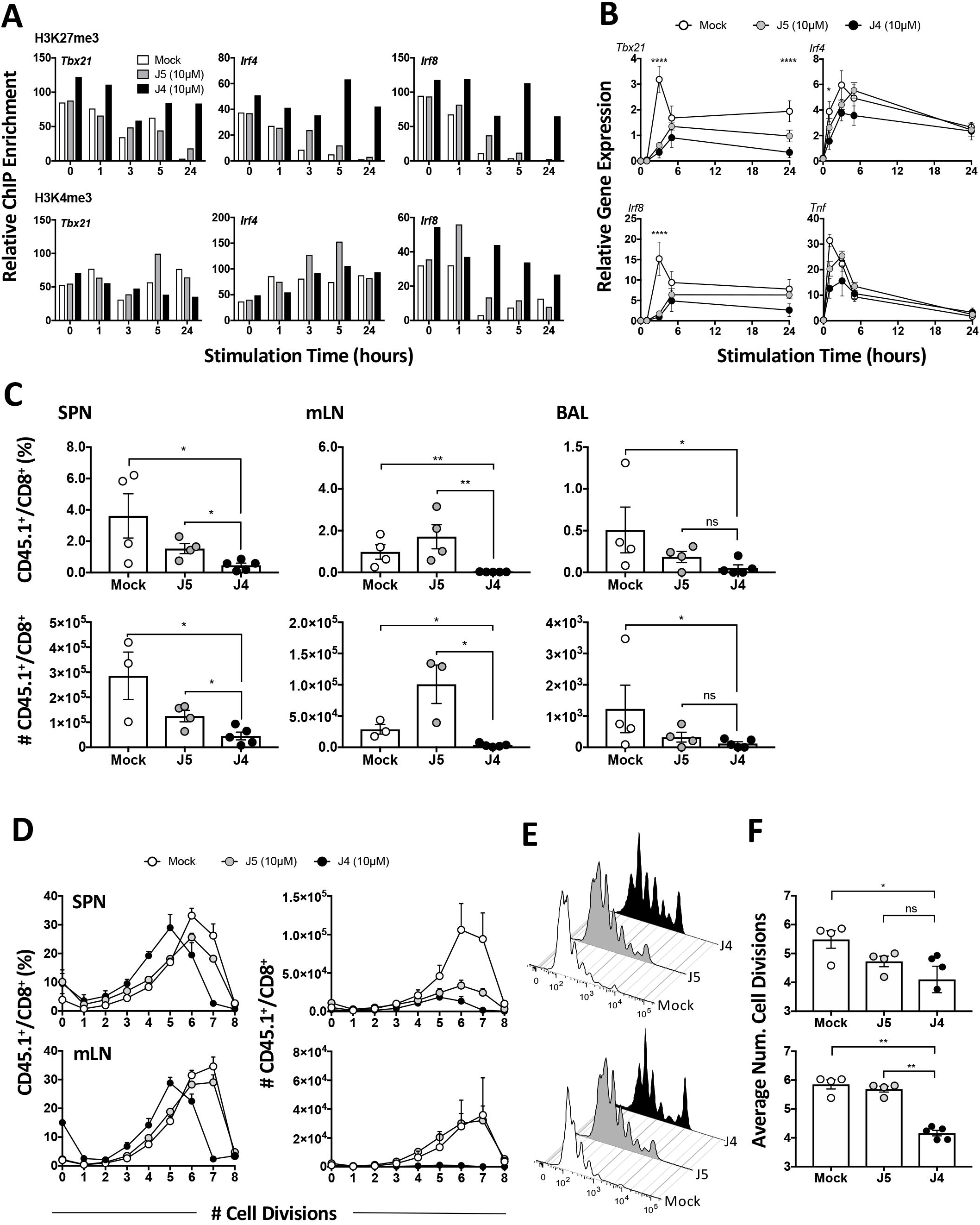
KDM6B inhibition prior to first cell division impairs CD8^+^ T cell expansion in response to activation. **A)** Sort purified OT-I naïve CD44^int/lo^ CD62L^hi^ CD8^+^ T cells were either left untreated (mock) or treated with 10μM of the J5 control and J4 inhibitor for 2hrs in the presence of IL-2 before activating with the N4 peptide for 0, 1, 3, 5 and 24 hours. Cells were then processed for ChIP-qPCR analysis for H3K27me3 and H3K4me3 at the promoters of *Tbx21, Irf4* and *Irf8*. Data is representative of 2 independent repeats. **B)** The same samples were also used to assess transcriptional regulation of *Tbx21, Irf4, Irf8* and *Tnf* using quantitative PCR. Data shown are mean ± SEM from 3 independent repeats. Statistical significance was calculated using 2 way-ANOVA (J4 vs. mock and/or J5 control, *p<0.05 **p<0.01, ***p<0.001). **C)** *In vivo* analysis of KDM6B inhibition on IAV-specific CD8^+^ T cell responses. CTV labelled OT-I CD8^+^ T cells were untreated (mock) or treated with GSK-J5 or GSK-J4 for 4hrs prior to adoptive transfer into C57BL/6 recipients, previously infected 3 days before with 10^4^ pfu X31-OVA. The proportion and number of CD45.1^+^/CD8^+^ OT-I T cells detected in the spleen (SPN), mediastinal lymph node (mLN) and the bronchoalveolar lavage (BAL) fluid were assessed by flow cytometry. **D)** The percentage and number of OT-I CD45.1^+^/CD8^+^ T cells undergoing different number of cell divisions was assessed. **E)** Representative FACS plot comparing the frequency of dividing OT-I CD45.1^+^/CD8^+^ T cells with mock, J5 or J4 treatment. **F)** The average number of cell divisions of CD45.1^+^ OT-I CD8^+^ T cells were compared between mock, J5 and J4 conditions. Data shown are mean ± SEM with 4-5 mice/group and are representative of 2 independent repeats. Statistical significance calculated using a two-tailed Student’s T-test (*p<0.05 **p<0.01, ***p<0.001).

Given the capacity of GSK-J4 to inhibit early H3K27 demethylation after CD8^+^ T cell activation, we next determined whether GSK-J4 treatment of naive OT-I CD8^+^ T cells would impact virus-specific CD8^+^ T cell differentiation. Naive, CellTrace Violet (CTV) labelled OT-I T cells were treated with the GSK-J4 inhibitor, or GSK-J5 analogue for 4hrs *in vitro*. An equal number of treated OT-I T cells were then adoptively transferred into recipient B6 mice infected 3 days prior to transfer with the A/HKx31-OVA virus (**Fig. 5C**). Pre-treating OT-I T cells with either the J5 control or the GSK-J4 inhibitor did not affect the viability of these cells nor the expression of CD44 or CD62L prior to transfer (**Supplementary Figure. S5**). At 3 days after transfer, both the proportion and absolute number of GSK-J4-treated OT-I CD8^+^ T cells were reduced compared to the mock or GSK-J5 treated OT-I CD8^+^ T cells (**Fig. 5C**). While both mock or J5-treated OT-I T cells had undergone extensive cell division (**Fig. 5D-F)** with an average of 5-6 cell divisions (**Fig. 5F**), GSK-J4-treated OT-I CD8^+^ T cells had undergone fewer divisions **(Fig. 5F)**. This result complimented our earlier bioinformatic analysis that indicated that H3K27me3 demethylation was required to engage gene networks involved in cell division and cell cycling (**Fig. 3D**). Therefore, the inability to efficiently demethylate H3K27me3 early after CD8^+^ T cell activation resulted in a diminished capacity to fully engage the proliferative capability of virus-specific CD8^+^ T cells in response to infection.

Examination of functional characteristics of the responding OT-I T cells demonstrated that early H3K27 methylation was also required for acquisition of lineage-specific functions. For example, GSK-J4 treated OT-I T cells had both a lower proportion of T-BET^+^ CD8^+^ T cells and expressed lower levels of T-BET and GATA3 within positive cells (**Supplementary Figure. S6A, B**). Interestingly, there was a lower proportion of GSK-J4 treated OT-I T cells located within the draining lymph node producing IL-2, IFN-γ or TNF upon reactivation, compared to mock treated and GSK-J5 treated cells (**Supplementary Figure S6C-E**). This was also reflected in a diminished proportion of multifunctional OT-I T cells isolated from the draining lymph node capable of simultaneously producing all three cytokines (**Supplementary Figure S6F-G**). Importantly, there was no difference in the amount of cytokine produced. These data support early reports showing that acquisition of CD8^+^ T cell effector function is linked to cellular division (Denton et al., 2011; Lawrence and Braciale, 2004). Hence, H3K27 demethylation, prior to initial cell division, appears to be a critical step for initiation of the autonomous CD8^+^ T cell differentiation program induced by T cell activation.

### H3K27me3 removal is required for establishing virus-specific CD8^+^ T cell memory

It has previously been demonstrated that diminished effector responses can still lead to effective CD8+ T cell memory populations (Badovinac et al., 2004; Zehn et al., 2009). We therefore next examined whether CD8+ T cell memory T cell formation was left intact after inhibition of KDM6B-dependent H3K27me3 demethylation prior to initial activation. Naive OT-I CD8^+^ T cells treated with either the GSK-J5 analogue, or the GSK-J4 drug, and were adoptively transferred into B6 recipient mice that had been infected with A/HKx31-OVA one day prior. The primary and secondary OT-I responses were then assessed at the peak of the primary response (day 10), memory (day 30) or after secondary challenge with A/PR8-OVA (day 6, secondary) (**Fig. 6A**). In support of our earlier data, GSK-J4 treatment had a profound impact on the expansion of OT-I CD8^+^ T cells optimal during the primary acute effector response. The diminished primary response observed after GSK-J4 treatment of OT-I CD8^+^ T cells was also reflected in the establishment of a lower memory OT-I cell frequency in the spleen (**Fig. 6B**). Comparison of OT-I CD8^+^ T cell numbers in the lung tissue 30 days after primary infection demonstrated that GSK-J4 treatment prior to adoptive transfer resulted in a significantly reduced frequency and number of total memory OT-I CD8^+^ T cells (**Fig. 6C**). Utilising intravital injection of anti-CD3 antibody to distinguish resident versus circulating memory CTL, we determined that GSK-J4 treatment also resulted in fewer tissue resident CD69^+^CD103^+^ memory OT-Is (**Fig. 6D**). Thus it appears that inhibition of H3K27me3 demethylation impacted the formation of effector and memory CTL populations.

**Figure 6.**
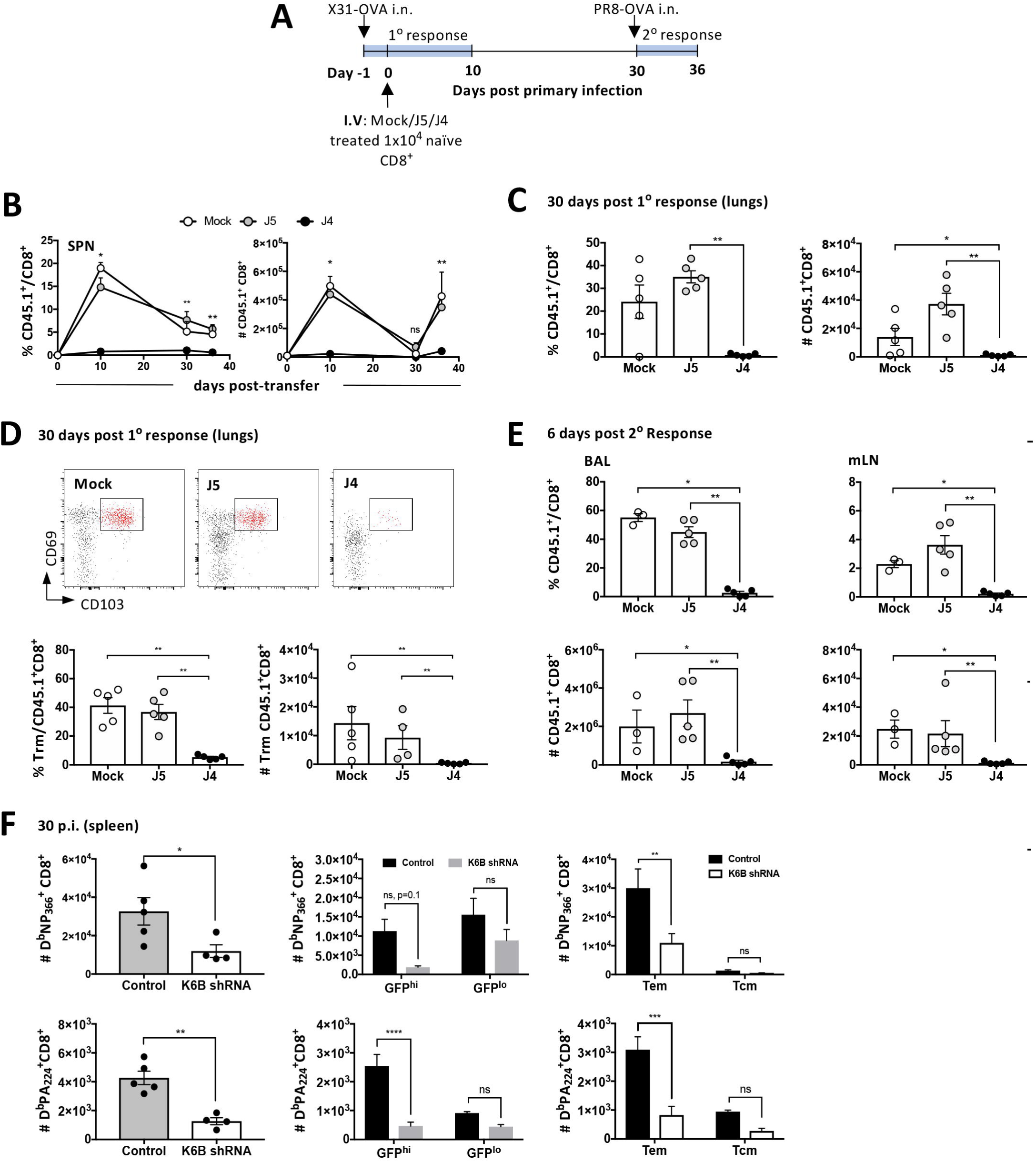
Antigen-specific memory formation requires H3K27me3 removal during early CD8^+^ T cell activation. **A)** OT-I CD8^+^ T cells that were either mock treated, or treated with GSK-J5 or GSK-J4 for 4hrs prior, were adoptively transferred into C57BL/6 recipients infected with 10^4^ pfu X31-OVA a day before (1° response). Mice were rested for 30 days before re-challenge with 10^4^ pfu PR8-OVA (2° response). **B)** The proportion and number of CD45.1^+^CD8^+^ OT-I T cells were enumerated in the spleen of mice on day 10 and 30 post-transfer and again 6 days post-secondary challenge with PR8-OVA. These were compared between the mock, J5 and J4 treatment. Data shown are mean ± SEM with 4-5 mice/group and are representative of 2 independent repeats. **C)** The proportion and the number of memory CD45.1^+^CD8^+^ OT-I T cells were enumerated in the lungs of mice 30 days post primary infection with X31-OVA. **D)** Representative FACS plot of CD45.1^+^CD8^+^ resident memory T cells (CD69^hi^CD103^hi^) identified in the lungs mice that received CD8^+^ OT-I T cells that received mock, GSK-J5 or GSK-J4 treatment. The proportion and number of mock, GSK-J5 or GSK-J4 treated CD45.1^+^CD8^+^ Trm cells were compared. Data shown are mean ± SEM with 4-5 mice/group and are representative of 2 independent repeats. **E)** The proportion and number of mock, GSK-J5 or GSK-J4 treated memory CD45.1^+^CD8^+^ OT-I T cells were enumerated in the BAL and mLN from mice that were challenged with PR8-OVA. Data shown are mean ± SEM with 4-5 mice/group and are representative of 2 independent repeats. **F)** The number of IAV-specific (D^b^PA_224_ and D^b^NP_366_) CD8^+^ T cells quantified in the spleen in VAV-tTA Lc1309 (control) and VAV-tTA Kdm6B shRNA mice 30 d.p.i. with 10^4^ pfu X31. The number of IAV-specific CD8^+^ T cells compared between GFP (shRNA) high versus low cells between the control and Kdm6b shRNA expressing mouse strains. The number of IAV-specific Tem and Tcm CD8^+^ T cells were compared between the control and Kdm6b shRNA expressing mouse strains. Data shown are mean ± SEM with 4-5 mice/group. All statistical significance shown here were calculated using a two-tailed Student’s T-test (*p<0.05 **p<0.01, ***p<0.001).

It has been previously demonstrated that limiting initial effector T cell expansion does not necessarily impact the recall capacity of memory CD8^+^ T cells (Zehn et al., 2009). To determine whether inhibiting H3K27me3 demethylation also impacted the recall capacity of established memory OT-I, we secondarily challenged primarily infected mice that had received GSK-J4-, J5- or mock treated OT-Is with a serologically distinct A/PR8 strain of IAV-OVA (**Fig. 6B, E**). Memory OT-I T cells established after transfer and primary activation of GSK-J4-treated OT-Is failed to expand upon secondary infection. This was evident in the spleen (**Fig. 6B**), the bronchoalveolar lavage fluid (BAL) and mLN (**Fig. 6E**). Together these data suggest that a failure to remove H3K27me3 early after activation not only impacts initial T cell expansion, but also programming of memory recall potential. This was likely not due to a difference in starting memory T cell number as the fold expansion in GSK-J4 treated mice was significantly diminished compared to the GSK-J5 controls. This indicates an intrinsic defect in recall capacity.

To further consolidate our finding that KDM6B plays a role in programming optimal CD8^+^ T cell memory, we utilised transgenic mice that constitutively express a *Kdm6b*-specific shRNA, for constitutive knockdown of *Kdm6b* (Prier et al., 2019a). The mice express *Kdm6b* mRNA as part of a GFP reporter where the levels of GFP are indicative of shRNA levels (Prier et al., 2019a). Expression of the *Kdm6b* shRNA knocked down *Kdm6b* transcription by ∼50% in naïve CD8^+^ T cells compared to the luciferase shRNA control (**Supplementary Figure S8**). *Kdm6b* shRNA mice were infected with A/HKx31 and memory D^b^NP_366_ and D^b^PA_224_-specific CD8^+^ T cell responses analysed in the spleen 30 days after infection. *Kdm6b* knockdown in GFP^hi^ CD8^+^ T cells resulted in a diminished number of IAV-specific memory CD8^+^ T cells compared to the luciferase shRNA knockdown controls (**Fig. 6F**). This diminished response was reflected in lower proportion of both effector (T_EM_) and central (T_CM_) memory subsets. Altogether, both of these experimental models demonstrate that inhibition of KDM6B, and subsequent H3K27me3 demethylation during early T cell activation, is critical for facilitating not just mature primary virus-specific CD8^+^ T cell expansion, but formation of functional virus-specific memory CD8^+^ T cells.

## DISCUSSION

Naive CD8^+^ T cell activation results in an autonomous program of cellular proliferation that results in the acquisition of lineage-specific function (Kaech and Ahmed, 2001; van Stipdonk et al., 2003). While the virus-specific CD8^+^ T cell differentiation process is highly controlled and regulated (Cruz-Guilloty et al., 2009; Intlekofer et al., 2005; Kallies et al., 2009; Marchingo et al., 2014; Schlub et al., 2009; Xin et al., 2016), while triggering CD8^+^ T cell proliferation only requires a short stimulus (Kaech and Ahmed, 2001), at least 20 hrs of stimulus is required for functional CD8^+^ T cell expansion. Our study provides molecular evidence that initial T cell activation results in rapid KDM6B dependent removal of H3K27me3 enabling engagement of transcriptional pathways required for preparing T cells for subsequent proliferation and differentiation. KDM6B-dependent removal of H3K27me3 enables increased chromatin accessibility and the staged exposure of specific TFBSs within gene regulatory elements. This is associated with subsequent histone acetylation and stable transmission of transcriptionally permissive chromatin structures into effector and memory CTL populations. An inability to appropriately engage these cellular support processes prior to first cell division negatively impacts subsequent optimal differentiation of virus-specific CTL and establishment of virus-specific CD8^+^ T cell memory.

It is now well accepted that genome wide changes in chromatin accessibility and post-translational histone modifications are associated with the transition of naive T cell differentiation into the effector/memory states (Denton et al., 2011; Northrop et al., 2008; Russ et al., 2014; Russ et al., 2017; Scott-Browne et al., 2016; Sen et al., 2016; Wei et al., 2009; Zediak et al., 2011). It has been demonstrated that dynamic modulation of H3K27me3 deposition is evident during thymic T cell development (Zhang et al., 2012). In this study, we demonstrated that TCR engagement specifically up-regulated *Kdm6b* transcription, and not *Kdm6a*, which is another H3K27me3 demethylase. This differential H3K27me3 demethylase upregulation after TCR activation potentially explains the observation that virus-specific effector and memory CD8^+^ T cell generation is normal in *Kdm6a* knockout mice (Cook et al., 2015; Yamada et al., 2019). Interestingly, T cell specific *Kdm6a*-deficiency did restrict LCMV-specific T follicular CD4^+^ T cell responses, leading to increased susceptibility (Cook et al., 2015). This may point to distinct roles for KDM6A and KDM6B in CD4^+^ and CD8^+^ T cell responses, respectively.

Interestingly, concurrent with KDM6B upregulation, we saw upregulation of the PRC2 H3K27me3 methyltransferase subunits, *Ezh2* and *Suz12*. This raises an interesting question about how recently activated CD8^+^ T cells initiate appropriate gene transcription when there is active competition between enzymes that write or erase H3K27me3. Our comprehensive profiling of H3K27me3 deposition in early-activated and *ex vivo* differentiated effector and memory T cells showed that regions that exhibited either delayed or stably-maintained H3K27me3 demethylation were associated with an initial permissive chromatin state that was consolidated by H3K27Ac^+^ enrichment and chromatin accessibility. Importantly, these regions exhibited significant overlap with publicly available ChIP-seq data identifying regions in activated T cells bound by the histone acetyltransferase, P300. In contrast, regions that showed transient H3K27me3 demethylation, overlapped with regions that were targets for the PRC2 subunit, SUZ12. It is tempting to speculate that a rapid transition from methylation to acetylation at H3K27 acts a molecular switch that ensures activation of stable gene transcription required for optimal T cell differentiation and proliferation. This then protects activated gene loci from the activity of opposing chromatin modifiers ensuring stability of differentiation state.

A recent study showed that EZH2 activity and H3K27me3 deposition, during the late effector T cell response, is required to repress pro-memory genes ensuring virus-specific CD8^+^ effector T cell generation (Gray et al., 2017). Interestingly, a similar study demonstrated that *Ezh2* upregulation within virus-specific CD8^+^ T cells that had undergone one cell division marked T cells committed to become effector T cells (Kakaradov et al., 2017). Our data suggest that such fate decisions may be made even prior to the first cell division as upregulation of *Ezh2* was observed at 3, 5 and 24hrs after activation.

Analysis of genomic regions that lost H3K27me3 and became more accessible over the time course, demonstrated staged exposure of specific TFBS before initial cell division. BATF/JUN binding sites emerged first, followed by STATs/IRF/NFAT/NFκB site, and finally TBX/RUNX sites at 24 hrs after activation. The initial unmasking of regions containing BATF/JUN family binding sites fits with earlier observations that BATF acts as an important factor for initiation of CD8^+^ T cell activation (Godec et al., 2015; Kurachi et al., 2014). Importantly, studies to date narrowed the timing of BATF activity to somewhere within the first 3 days after CD8^+^ T cell activation (Godec et al., 2015). Our data demonstrate that BATF activity is likely required in the very initial stages of T cell activation, and when paired with its binding partners (such as IRF4), it could potentially act as a pioneering factor helping remodel the chromatin landscape within hours of activation. Whether KDM6B recruitment is dependent on BATF, or if H3K27me3 removal at target gene loci precedes BATF binding to initiate T cell activation will be of interest to examine in the future.

KDM6B dependent removal of H3K27me3 was evident at the *Tbx21* locus as early as 3hrs, and stable up to 24hrs after activation. This correlated with rapid upregulation of *Tbx21* transcription prior to cell division. T-BET upregulation was prior to the emergence of TBX21 binding sites within H3K27 demethylated regions found at 24 hours after activation. We have recently shown that T-BET deficiency results in early dysregulation of virus-specific CD8^+^ T cell differentiation that results in an inability to expand (Prier et al., 2019b). This was associated with decreased H3 acetylation at a T-BET target transcriptional enhancer within the *Ifng* locus. It has been previously demonstrated that T-BET can physically interact with, and recruit H3K27 demethylases to the *Ifng* regulatory elements in CD4^+^ T_H_1 cells (Miller et al., 2008). Hence, we hypothesise that T-BET works downstream of pioneering factors such as BATF/IRF4/RUNX3, by targeting CD8^+^ T cell specific gene loci to further modulate chromatin remodelling. These results are reminiscent of a mechanism observed in stem cell differentiation where early H3K27me3 removal, after receipt of differentiation signals, enables binding of lineage-specifying TFs to newly remodelled chromatin structures and commitment to a differentiated cell fate (Agger et al., 2007).

Inhibition of KDM6B activity prior to T cell activation had a profound impact on subsequent virus-specific CD8^+^ T cell proliferation, and the capacity to establish an effector memory T cell pool. Naive CD8^+^ OT-I T cell treatment with the KDM6B inhibitor prior to adoptive transfer severely impacted the proliferative capacity of OT-I T cells responding to IAV infection. Despite the drug likely being diluted upon subsequent cell division, GSK-J4 treated OT-I CD8^+^ T cells exhibited delayed division kinetics and failed to fully expand to levels observed in control treated cells. This suggests that early H3K27me3 removal is of critical importance for subsequent clonal expansion and differentiation. Concomitant with diminished proliferation in the lymph node, we observed that GSK-J4 OT-I CD8^+^ T cells exhibited lower levels of T-BET expression and a lower proportion expressing multiple cytokines. These data further support the idea that molecular re-programming in the lymph node during the early stages of T cell activation is a key step for optimal effector T cell differentiation.

Deposition of H3K27me3 at key pro-memory genes has been reported to be important for the formation of optimal effector virus-specific T cell responses (Gray et al., 2017; Kakaradov et al., 2017). Similarly, it has also been recently shown that deposition of another repressive histone mark, H3K9me3, was required for shutting down pro-memory gene loci enabling optimal effector CD8+ T cell differentiation (Pace et al., 2018). Further, limiting T cell proliferation has also been observed to promote formation of memory CD8^+^ T cell populations (Badovinac et al., 2005; Badovinac et al., 2004; Zehn et al., 2009). This is supported by the notion that effector CTL are more terminally differentiated compared to memory CTL that maintain self-renewal capacity (Crompton et al., 2016). Hence, it was possible that inhibition of H3K27me3 removal would help promote memory formation. GSK-J4 treated OT-I CD8^+^ T cells exhibited an intrinsic failure to expand upon secondary IAV challenge. In particular, we observed profound defects in the formation of lung resident CD8^+^ memory OT-Is (T_RM_). Hence, KDM6B dependent H3K27me3 demethylation during the early stages of a primary T cell response impacts efficient programming of both effector and memory CD8^+^ T cell fates. This data also suggests that commitment to effector and memory T cell fates are independent processes. This might reflect the role of distinct TFs. For example, we observed sequential unmasking of specific TF motifs for EOMES and RUNX3 TFs that have been shown to alter CD8^+^ T cell differential potential for the formation of distinct memory T cell subsets (Mackay et al., 2015; Miller et al., 2008; Milner et al., 2017; Wang et al., 2018). By regulating the accessibility of these T cell lineage TF motifs during early hours of T cell activation, H3K27me3 demethylation therefore regulates the timely commitment to both effector and memory T cell fates.

## Acknowledgements

This work was supported by grants from the National Health and Medical Research Council of Australia (Program Grant #5671222 awarded to awarded to SJT and NLL; Project grant #APP1003131 awarded to S.J.T); and an Australian Research Council Discovery Grant (DP DP170102020 awarded to S.J.T and S.R); S.J.T is supported by an NHMRC Principal Research Fellowship; NLL is supported by an Australian Research Council Future Fellowship. We thank the Monash Genomics Platform (Micromon) for high throughput sequencing and the Monash Bioinformatics Platform for data analysis.

## Author Contributions

Conceptualisation, S.J.T, J.L. and S.R. Methodology, J.L, K. H, M.O., L.J.G, X.Y.X.S, and A.B. Formal analysis, J.L., K.H., M.O., L.J.G., and X.Y.X.S. Investigation, J.L, J.E.P, X.Y.X.S., M.L.T.N., D.P., and B.R. Resources, S.J.T, S.R and P.J.H. Data Curation, K. H., M.O. and L.J.G. Writing original draft, S.J.T and J.L; Writing-Review and Editing, S.J.T, J.L., N.L.L, K.H., B.R. and S.R. Supervision, S.J.T. Acquisition of funding, S.J.T., P. J. H., N.L.G and S.R.

## Declaration of interests

S.R is currently the Founder/Chief Scientific Officer of EpiAxis Therapeutics. This manuscript received no funding from EpiAxis for this work. S.J.T is a member of the scientific advisory board for Medicago Inc, Quebec, Canada. This manuscript received no funding from Medicago for this work.

## STAR METHODS

### Cell preparation

Naïve OT-I CD8^+^CD44^lo/int^ cells were purified from OT-I/Ly5.1 male mice (6-8 weeks) (>99% purity). They were stimulated with 1μM OVA (N4) for 0, 1, 3, 5 and 24 hours at the presence of rhIL-2 (10U/mL) in cRMPI. For demethylase inhibition, purified OT-I CD8^+^ T cells were pre-treated with 10μM of the control inhibitor (J5) or the Kdm6b inhibitor (J4) for 2hrs in cRPMI with rhIL-2 (10U/mL) before stimulation with the N4 peptide for 0, 1, 3, 5 and 24 hours.

### In Vivo Histone Demethylase Inhibition

Total lymph nodes were extracted from OT-I females (6-10 weeks). They were resuspended to generate a single cell suspension followed by labelling with the CellTrace Violet Cell Proliferation kit. A total of 4×10^6^ lymphocytes were used for either the mock treatment or pretreatment with either the substrate specificity control (J5) or the H3K27me3 demethylase inhibitor (J4) in rhIL-2 (10U/mL) for 4hrs. A portion of these cells were used to stain with the Annexin V kit with PI and anti-CD44-PE-Cy7, CD62L-BV570, CD8-APC and CD45.1-PE antibodies. A proportion of 3×10^5^ naïve (CD44^int/lo^CD62L^hi^) CD8^+^ cells were then intravenously injected into female C57BL/6 mice (6-8 weeks) that had been infected with 10^4^pfu x31-OVA for 3 days (early time point) or 10^4^ naïve (CD44^int/lo^CD62L^hi^) CD8^+^ cells were used for determining early memory formation at day 30 post-infection. At these time points, the spleen (SPN), mediastinal lymph nodes (mLN)and bronchoalvelolar lavage (BAL) fluid or the lungs were extracted to prepare for single cell suspension. This was followed by staining with the live/dead aqua-blue dye, anti-CD45.1, CD45.2, CD8, Gata3, T-bet, IFN-g, IL-2, TNF antibodies for flow cytometric analysis.

### Flow Cytometry

For flow cytometry analysis, antibody-stained samples were acquired on then FACSCanto II or the Fortessa flow cytometers (BD Biosciences) coupled to the high through system (HTS). Post-acquisition data analyses were performed using FlowJo software (Tree Star, Ashland, OR, USA). Mean fluorescence intensity (MFI) or the frequency (%) of staining was determined as the geometric mean of positive population.

### Total RNA extraction

Total RNA was extracted using Trizol® from unstimulated or stimulated OT-I CD8^+^ T cells. For gene expression analysis, 100μg mRNA was converted to cDNA using the Omniscript kit (Invitrogen) according to manufacturer’s instructions. Relative gene expression changes were determined by quantitative real time-PCR using the CFX-Connect Real-Time System (Biorad) with Taqman ® Gene MGB primer/probes (Life Technologies).

### RNA-sequencing

RNA samples (triplicates) were depleted of DNA and purified using the Qiagen RNeasy MinElute kit. The bioanalyzer was used to determine the integrity of the RNA before library preparation using the kit. RNA libraries were sequenced paired end (100bp) on the Hiseq2000 instrument at the Australian Genome Research Facility, the Walter and Eliza Hall Institute of Medical Research, Melbourne, Australia. Data quality was confirmed with fastqc software. Paired end RNA-seq data was mapped to mouse genome mm10 using TopHat (with Bowtie2). Only concordant pairs with mapping quality greater than 10 were utilised. Reads were assigned to annotated genes using Feature Counts from R subread Bioconductor R package. Genes which did not have at least 3 counts in each sample in at least one group were excluded from the analysis. Differential Expression analysis was done using edgeR Bioconductor R package. Genes were considered DE if exhibited FDR < 0.05 and log2 FC > 1. Log averages of the triplicates for the differential genes were clustered using Manhattan distance, complete linkage (R) and grouped using the z-score of the averaged TMM normalized values (EdgeR) with K means (R).

### Chromatin Immuno-precipitation (ChIP) and Formaldehyde-assisted Isolation of Regulatory Element (FAIRE) Assays

Cells crosslinked with 0.6% formaldehyde were sonicated and immune-precipitated with anti-H3K4me3 and H3K27me3 ChIP-grade antibodies and Protein A magnetic beads (Millipore). Total input and no-antibody control for each sample was included for normalisation and specificity control purposes. Immuno-precipitated DNA was purified and re-suspended in 0.1X TE buffer. For FAIRE-analysis, open chromatin was extracted twice by adding an equal volume of phenol:chloroform:isoamyl (25:24:1) (Sigma) and precipitated as described for ChIP assays. Resulting ChIP or FAIRE-DNA was compared using quantitative real time-PCR on the CFX-Connect Real-Time System (Biorad) with Sybr-green master mix using primers spanning region of interest. Real-time PCR cycle threshold (Ct) values were converted to copy number and background immunoprecipitation subtracted (no-antibody control).

### ChIP-sequencing

ChIP-DNA was prepared for sequencing using the NEBNext® CHIP-seq Library Prep Master Mix Set for Illumina (NEB #E6240L, New England BioLabs Inc) according to manufacturer’s instructions. ChIP-DNA library was subjected to size selection with AMPure beads. The quality of the ChIP-DNA library and the fragment size of approximately 275 bp were assessed on the Bioanalyzer using the Agilent High Sensitivity DNA chip (Agilent Technologies, 5067-4626). ChIP libraries were sequenced paired end on the Hiseq2500 instrument at the Australian Genome Research Facility (AGRF). Differential ChIP-seq peaks were found by creating windows of counts (bigwig files normalized per 10 million reads (HOMER)) for each treatment, finding the differences between windowed counts (DeepTools) and then calling peaks in MACS2 (bdgpeakcall -c 3 -l 150 -g 300) (Zhang et al., 2008). Regions were intersected using Bedtools and annotated using HOMER, CISTROME (Liu et al., 2011) (in conjunction with BedTools) and transcription start sites taken from ENSEMBL. DAVID and Metascape (Zhou et al., 2019) were used to examine gene groups for enriched Gene Ontology terms.

### ATAC-seq

ATAC-seq is adapted from (Buenrostro et al., 2015). A total of 50 000 cells were lysed with cold lysis buffer for nuclei extraction. Nuclei were immediately resuspended in transposition reaction mix prepared from the Illumina Nextera DNA Sample Preparation Kit (FC-121-1030) for 30 minutes at 37°C. Transposed DNA was extracted using the Qiagen MinElute PCR Purification kit (Cat #. 28004). Resulting DNA was subjected to 5 PCR cycles on the thermocycler using a PCR primer 1 (Ad1_noMX) and an indexed PCR primer 2. An aliquot of each sample was used subsequently in a real-time quantitative PCR for 20 cycles to determine the number of cycles required for library amplification. The amplified DNA was purified using the Qiagen MinElute PCR Purification kit. Library quality was assessed using the bioanalyzer (Agilent) to ensure that the DNA fragmentation ranges between 50-200bp and the Qubit to determine the overall DNA concentration. ATAC-DNA was sequenced paired end on the Hiseq2500 instrument at the Australian Genome Research Facility (AGRF).

### Ciiider Analysis

CiiiDER analyisis was carried out according to (Gearing et al., 2019; Russ et al., 2017). Briefly, peaks with a length greater than 400 were filtered out. Regions of equal size were defined from 200 bases upstream to 200 bases downstream of the middle of each peak using the Mus_musculus.GRCm38.dna.primary assembly.fa genome. CiiiDER analysis was performed on these regions with JASPAR_CORE_vertebrates_2016.txt transcription factor position frequency matrices and a deficit cut-off of 0.15.

