## Supplementary figures for "KDM6B-dependent chromatin remodelling underpins effective virus-specific CD8^+^ T cell differentiation"

### Supplemental Figure 1

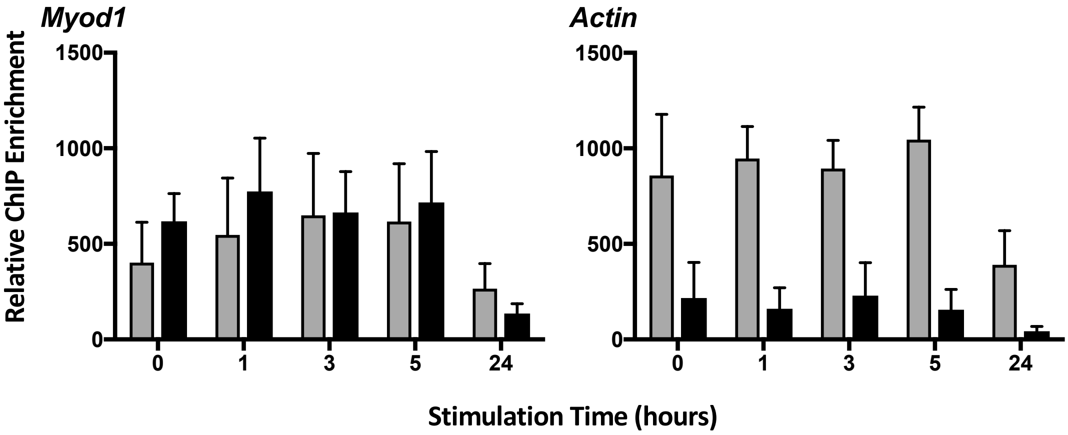


**Supplemental Figure S1. H3K4me3 and H3K27me3 ChIP for *MyoD* and *Actin*.** H3K27me3 (black bars) and H3K4me3 (grey bars) enrichment was measured by ChIP-qPCR on either naive OT-I CD8+ T cells or OT-I CD8+ T cells stimulated for 1, 3, 5 and 24hrs with N4 peptide. Enrichment was detected by quantitative PCR using primers spanning the promoter of *MyoD* and *Actin* promoters. Data shown are mean ± SEM with 3-4 independent samples.

### Supplemental Figure 2

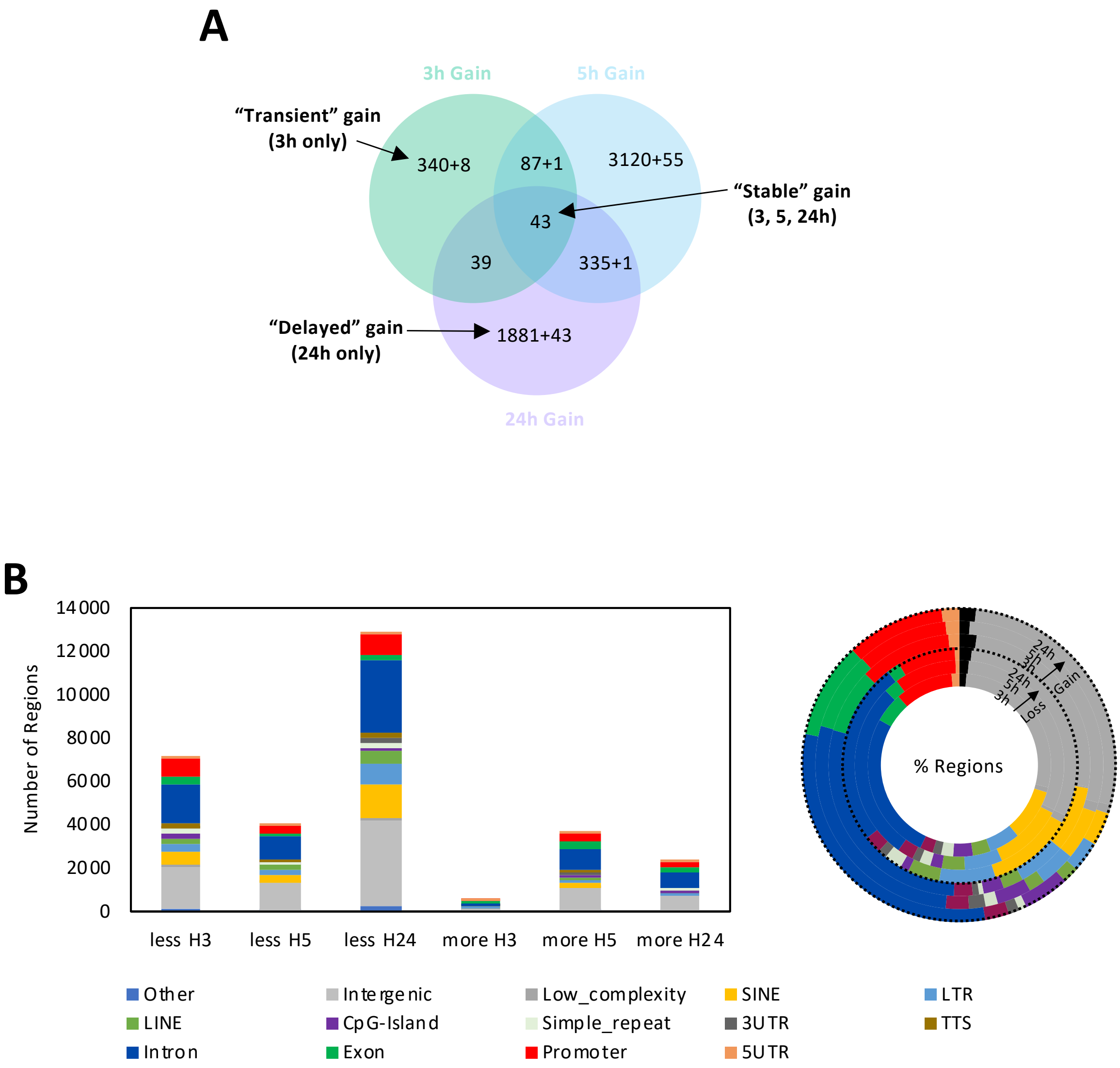


**Supplemental Figure S2. Genomic annotation of H3K27me3 regulated regions. A)** H3K27me3 ChIP-seq was performed on either naive CD8$^+$ OT-I T cells, or on OT-I T cells after 3, 5, or 24 hrs of *in vitro* stimulation as described above in Figure 1. Data was mapped back to the mouse genome (version mm10). Genomic regions that gained H3K27me3 within CD8$^+$ OT-I T cells activated for 3, 5 and 24 hours were compared to the unstimulated sample. These regions were categorised into either "transient", "delayed" or "stable" gain of H3K27me3 and enumerated. **B)** Using H3K27me3 ChIP-seq data generated above, regions that exhibited either H3K27me3 loss or gain were mapped to specific genomic functional regions. Data shown is the number and the proportion of regions displaying either H3K27me3 loss or gain in OT-I CD8$^+$ T activated for 3, 5 and 24hrs and compared to the unstimulated sample

### Supplementary Fig. 3

A

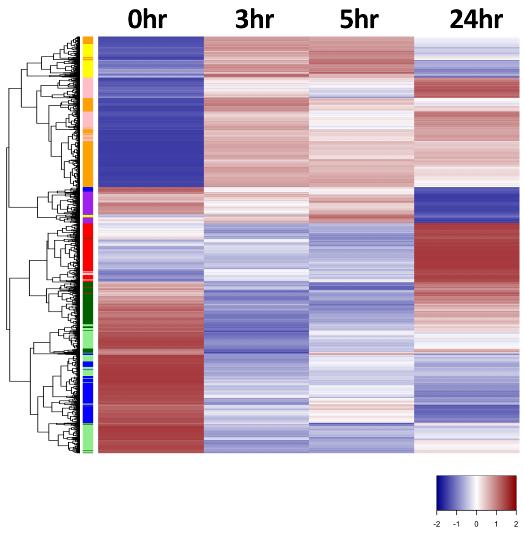


B

Transcriptional Modules

| T= | 0 | 3 | 5 | 24 | Gene Number |
|---|---|---|---|---|---|
| a | - | ↑↑ | ↑↑ | - | 491 |
| b | - | ↑ | ↑ | ↑ | 1124 |
| c | - | ↑ | ↑ | ↑↑ | 689 |
| d | - | - | - | ↑↑ | 798 |
| e | - | ↓↓ | ↓↓ | - | 747 |
| f | - | ↓ | ↓ | ↓ | 1001 |
| g | - | ↓ | ↓ | ↓↓ | 823 |
| h | - | - | - | ↓↓ | 415 |

C

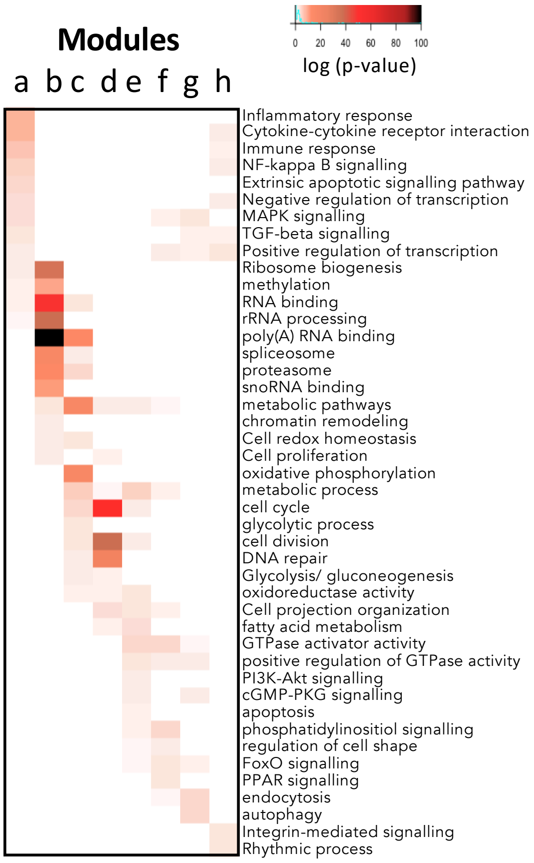


**Supplementary Figure S3. Naive T cell activation results in step-wise engagement of distinct transcriptional modules during the early hours of T cell activation. A)** *In vitro* activated OT-I $CD8^{+}$ T cells were subjected to RNAseq analysis as described in Figure 1. Significantly differentially expressed genes (DEGs) were clustered via hierarchical clustering (Manhattan) of z-scores based on $Log_2$ averages of triplicates and annotation of the k-means cluster on the left. **B)** Genes were categorised into distinct transcriptional modules (a – h) based on the pattern of up-regulation or down regulation at various times after initial activation, with the gene number for each module displayed for each group. **C)** Ontological analysis of the transcriptional modules was carried for DEGs either up or down regulated. Data shows the $log_2$ p-value for enrichment of genes enriched for specific ontologies for each module.

### Supplemental Figure 4

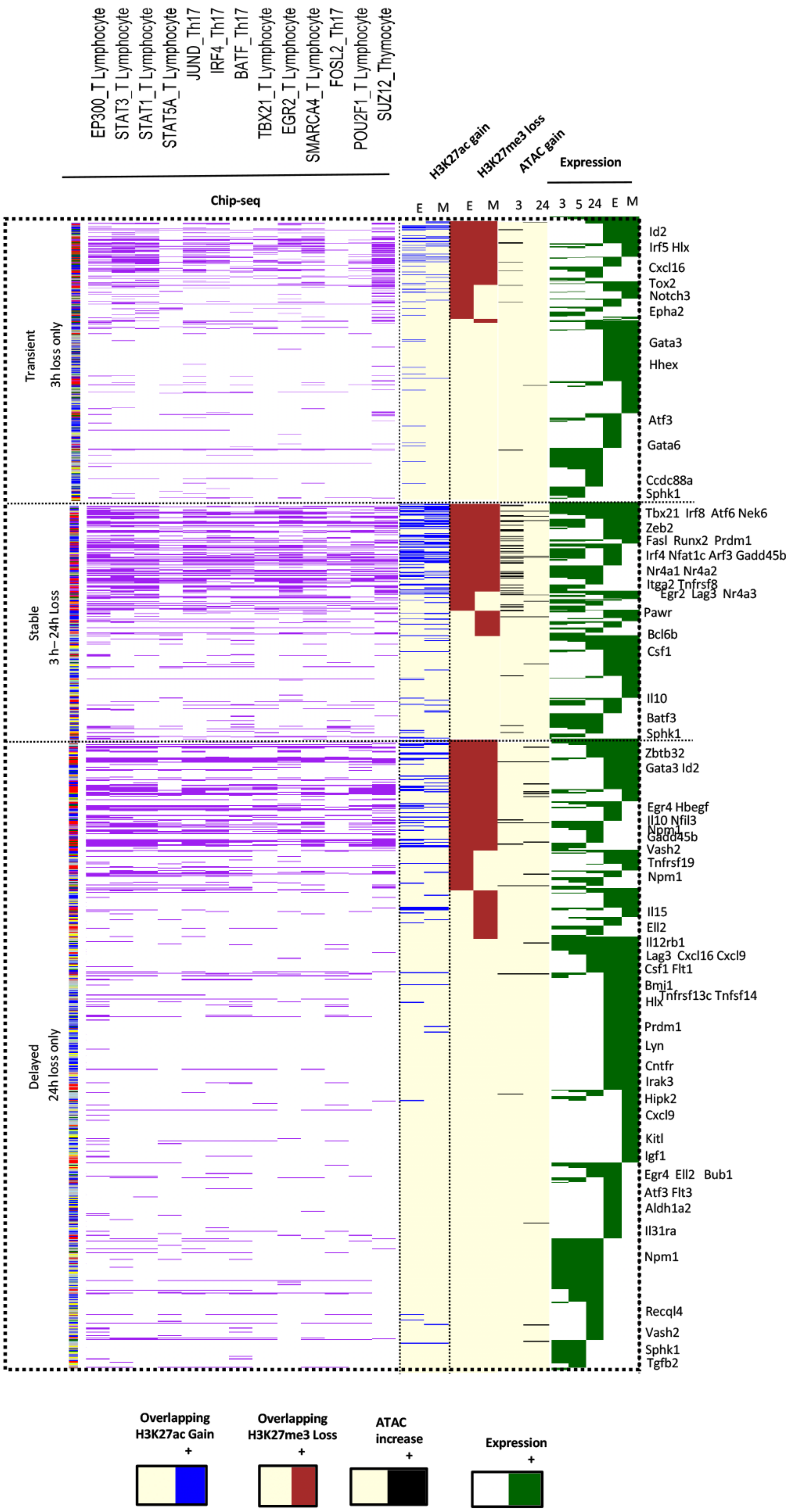

**Supplementary Figure S4. Stable H3K27me3 demethylation is associated with establishment of permissive epigenetic signatures and evidence of enriched TF binding.** H3K27me3 demethylated regions that were transient, stable or delayed in the demethylation patters were annotated to induced genes in unstimulated (0hr), early-activated OT-I CD8$^{+}$ T cells (3, 5, 24), *ex vivo*-derived OT-I effector (E) and memory (M) T cells. Regions that also overlapped published transcription factor bound regions are annotated in purple. Regions overlapping those exhibiting gain of H3K27ac gains (blue), and chromatin accessibility (ATAC) gain at 3 or 24hrs (black) were annotated. The genomic annotation of the region is shown on the right.

### Supplemental Figure 5

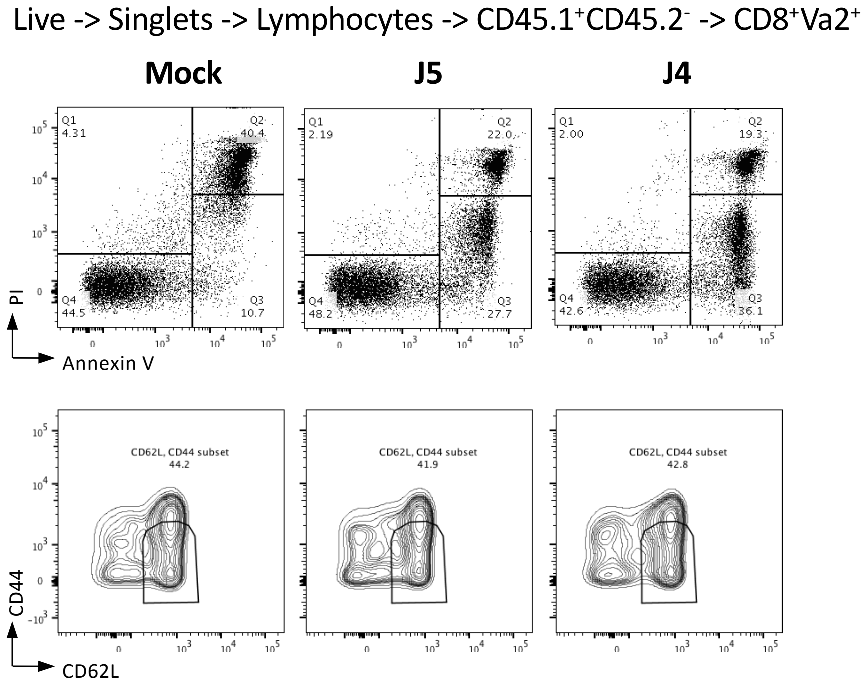


**Supplementary Figure S5. Viability of OT-I CD8$^+$ T cells after GSK-J4 treatment.** Naive CD8$^+$ OT-I lymphocytes were either untreated (mock) or treated with 10μM GSK-J5 control or GSK-J4 inhibitor for 4 hours *in vitro*. The viability of these OT-I CD8$^+$Va2$^+$ T cells was assessed using Annexin V and PI staining before adoptive transfer while the proportion of naïve CD8$^+$ T cells was measured between these treatments by measuring the expression of CD44 and CD62L. Representative flow cytometry plots are shown.

### Supplemental Figure 6

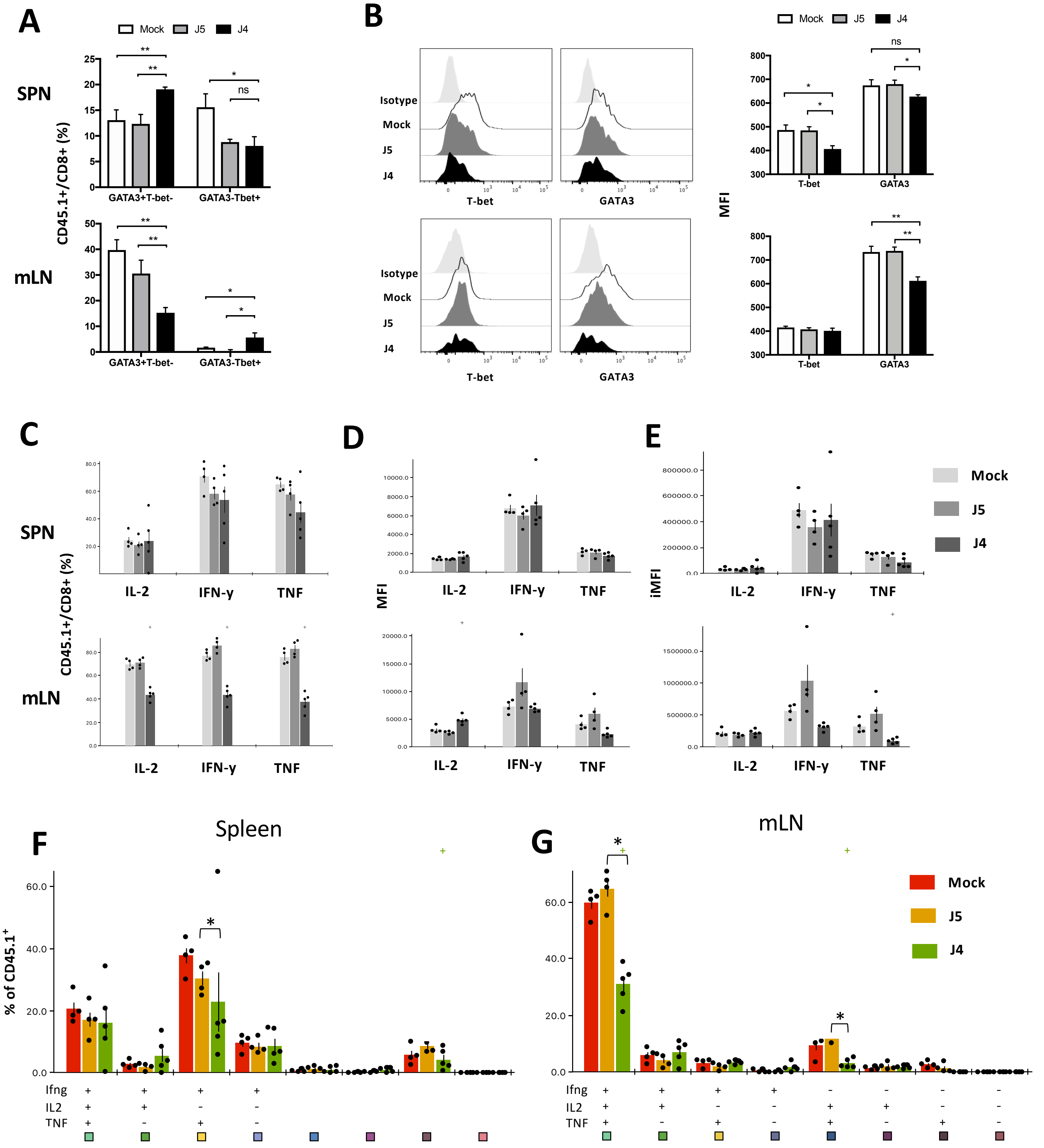

**Supplementary Figure S6. KDM6B inhibition prior to cell activation results diminished effector function within virus-specific CD8$^+$ T cells. A)** OT-I lymphocytes were either untreated (mock) or treated with 10μM GSK-J5 control or GSK-J4 inhibitor for 4 hours *in vitro*. 3x10$^5$ naïve (CD44$^{int/lo}$CD62L$^{hi}$) CD8$^+$ cells were then adoptively transferred into C57BL/6 mice infected with 10$^4$ pfu x31-OVA 3 days prior. Two and half days after transfer, percentage and number of GATA3$^{+/-}$T-BET$^{+/-}$ CD45.1$^+$/CD8$^+$ T cells were enumerated in the spleen (SPN) and mediastinal lymph node (mLN) across the different treatments. **B)** Data show representative histograms and Mean Fluorescence Intensity (MFI) of T-BET and GATA3 expression within splenic CD8$^+$ OT-I T cells that were mock treated, or treated with GSK-J5 or J4 prior to transfer was assessed. **C)** The capacity of mock treated, or GSK-J5 or J4 treated CD8$^+$ OT-I T cells to express cytokines was assessed by intracellular cytokine staining. Shown is the proportion of CD45.1$^+$/CD8$^+$ OT-I T cells expressing IL-2, IFN-γ and TNF, **D) t**he MFI of cytokine expression, **E)** total amount of cytokine produced, and F-G) the polyfunctionality of CD8$^+$ OT-I T cells was assessed for the spleen (SPN) and mediastinal lymph node (mLN) in the recipient mice across the different treatments. Data shown are mean ± SEM with 4-5 mice/group are representative of 2 independent repeats. Statistical significance calculated using a one-tailed Student's T-test (*p<0.05 **p<0.01, ***p<0.001).

### Supplemental Figure 7

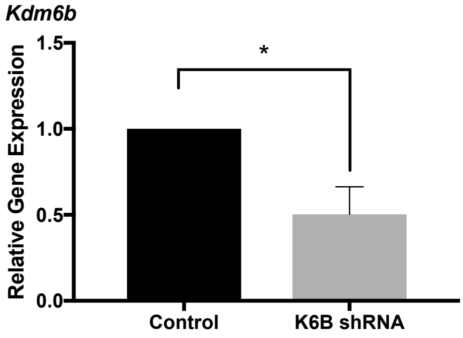


**Supplementary Figure S7. Confirmation of Kdm6b down-regulation in the VAV-tTA Kdm6b shRNA mouse.** Relative gene transcription of *Kdm6b* in naïve ($CD44^{int/lo}CD62L^{hi}$) $CD8^{+}$ T cells isolated from VAV-tTA Lc1309 (control) and VAV-tTA Kdm6B shRNA (K6B shRNA) was assessed by quantitative PCR using *Kdm6b* primer probes. Data shown is represented as mean ± SEM from 3 independent repeats with statistical significance calculated using a two-tailed Student's T-test (*p<0.05 **p<0.01, ***p<0.001).
